# Faecal virome transplantation decrease symptoms of type-2-diabetes and obesity in a murine model

**DOI:** 10.1101/792556

**Authors:** Torben Sølbeck Rasmussen, Caroline M. Junker Mentzel, Witold Kot, Josué L. Castro-Mejía, Simone Zuffa, Jonathan Swann, Lars Hestbjerg Hansen, Finn Kvist Vogensen, Axel Kornerup Hansen, Dennis Sandris Nielsen

## Abstract

**Objective:** Development of obesity and type-2-diabetes (T2D) are associated with gut microbiota (GM) changes. The gut viral community is predominated by bacteriophages (phages), which are viruses that attack bacteria in a host-specific manner. As a proof-of-concept we demonstrate the efficacy of faecal virome transplantation (FVT) from lean donors for shifting the phenotype of obese mice into closer resemblance of lean mice.

**Design:** The FVT consisted of viromes extracted from the caecal content of mice fed a low-fat (LF) diet for fourteen weeks. Male C57BL/6NTac mice were divided into five groups: LF (as control), high-fat diet (HF), HF+Ampicillin (Amp), HF+Amp+FVT and HF+FVT. At week six and seven of the study the HF+FVT and HF+Amp+FVT mice were treated with FVT by oral gavage. The Amp groups were treated with ampicillin 24 h prior to first FVT treatment.

**Results:** Six weeks after first FVT the HF+FVT mice showed a significant decrease in weight gain compared to the HF group. Further, glucose tolerance was comparable between the lean LF and HF+FVT mice, while the other HF groups all had impaired glucose tolerance. These observations were supported by significant shifts in GM composition, blood plasma metabolome, and expression levels of genes involved in obesity and T2D development.

**Conclusions:** Transfer of gut viral communities from mice with a lean phenotype into those with an obese phenotype reduce weight gain and normalise blood glucose parameters relative to lean mice. We hypothesise that this effect is mediated via FVT-induced GM changes.

**Significance of this study:** What is already known about this subject?

- Obesity and type-2-diabetes (T2D) are associated with gut microbiota (GM) dysbiosis.
- Faecal microbiota transplant from lean donors has previously shown potential in treating obesity and T2D.
- Patients suffering from recurrent *Clostridium difficile* infections (rCDI) have been cured with sterile filtered donor faeces (containing enteric viruses and no bacteria), here defined as faecal virome transplantation (FVT).

What are the new findings?

- FVT from lean donors lead to decreased weight gain and normalised blood sugar tolerance in a diet-induced obesity (DIO) mouse model.
- FVT significantly changed the bacterial and viral GM component, as well as the plasma metabolome and the expression profiles of obesity and T2D associated genes.
- Initial treatment with ampicillin prior FVT seems to counteract the beneficial effects associated with FVT.

How might it impact on clinical practice in the foreseeable future?

- We here show a proof-of-concept, providing a solid base for designing a clinical study of FVT targeting obesity and T2D in humans. This is further augmented by the increased safety related to FVT, since bacteria and other microorganisms are removed from the donor faeces, and therefore minimises the risk of disease transmission.
- These findings highlight the potential application of FVT treatment of various diseases related to GM dysbiosis and further support the vital role of the viral community in maintaining and shaping the GM.

## INTRODUCTION

Obesity and type-2-diabetes (T2D) constitute a world-wide health threat[1]. During the last decade it has become evident that certain diseases, including obesity and T2D, are associated with gut microbiota (GM) dysbiosis[2]. Interestingly, germ free (GF) mice do not develop diet-induced-obesity (DIO)[3], but when exposed to faecal microbiota transplantation (FMT) from an obese human donor they increase their body weight significantly more compared to GF mice exposed to FMT from a lean donor[4]. At present FMT is widely used to efficiently treat recurrent *Clostridium difficile* infections (rCDI)[5], and is suggested to have therapeutic potential against metabolic syndrome, a condition related to obesity and T2D[6]. FMT is considered as a safe treatment, however, safety issues remain since screening methods cannot fully prevent adverse effects caused by disease transmission from the donor faeces[7]. A failure in the screening procedure caused recently in a death (June 2019) of a patient following FMT[8] due to a bacterial infection.

The gut viral community (virome) is dominated by prokaryotic viruses[9], also called bacteriophages (phages), which are viruses that attack bacteria in a host-specific manner[10]. During recent years, evidence has mounted that the gut virome plays a key role in shaping the composition of the GM[11,12] as well as influencing the host metabolome[13]. Interestingly, rCDI have been successfully treated with a transfer of filtered donor faeces (containing phages, but no intact bacterial cells)[14]. Five out of five patients were successfully treated[14]. This approach is referred to as faecal virome transplantation (FVT). Further, antibiotic treatment alters the GM composition, but Draper *et al.* 2019 recently showed that the murine GM can be reshaped with FVT after antibiotic treatment[15].

DIO mice is a common animal model of metabolic syndrome, including symptoms such as obesity and insulin resistance/pre-T2D[16]. With C57BL/6NTac mice as the model, we hypothesised that FVT (originating from lean healthy donor mice) treatment of DIO mice would change the GM composition and directly or indirectly counteract the symptoms of obesity and T2D. To the best of our knowledge, this is the first study investigating the effect of FVT targeting obesity and T2D.

## METHODS

Detailed methods are enclosed in the online supplementary materials.

### Animal study design

Forty male C57BL/6NTac mice (Taconic Biosciences A/S, Lille Skensved, Denmark) were divided into 5 groups at 5 weeks of age: low-fat diet (LF, as lean control), high-fat diet (HF), HF+Ampicillin (Amp), HF+Amp+faecal virome transplantation (FVT) and HF+FVT (Figure 1). For thirteen weeks mice were fed *ad libitum* a HF diet (Research Diets D12492, USA) or a LF diet (Research Diets D12450J, USA). After six weeks on their respective diets, the HF+FVT and HF+Amp+FVT mice were treated twice with 0.15 mL FVT by oral gavage with a one-week interval (week 6 and 7) between the FVT. On the day before the first FVT inoculation the HF+Amp and HF+Amp+FVT animals were treated with a single dose of ampicillin (1 g/L) in the drinking water. The FVT viromes were extracted from the caecal content of eighteen mice fed a LF diet for 14 weeks, that were earlier isolated, sequenced, and analysed[17].

**Figure 1:**
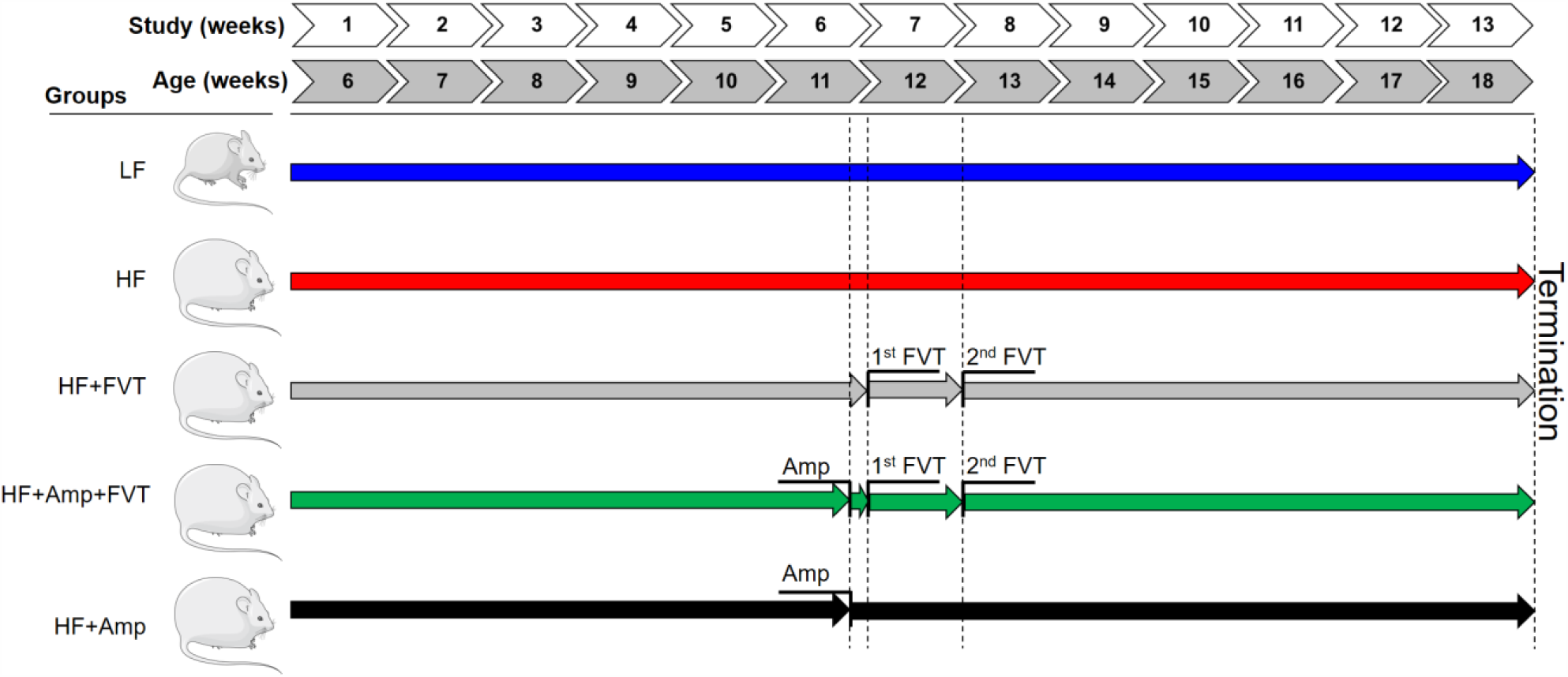
Experimental setup of the faecal virome transplantation (FVT). Forty male C57BL/6NTac mice (5 weeks old) were divided into five groups: Low-fat diet (LF, as healthy control), High-fat diet (HF), HF+Ampicillin (Amp), HF+Amp+FVT and HF+FVT. Their respective diets were provided continuously for 13 weeks until termination (18 weeks old). The HF+FVT and HF+Amp+FVT mice were treated with FVT twice with one-week interval by oral gavage at week 6 and 7 of the study. Ampicillin was added once to the drinking water one day before first FVT (week 6) for HF+Amp and HF+Amp+FVT. Mouse icon originates from https://smart.servier.com/under the CC BY 3.0 license.

The titer of the FVT viromes was approximately 2 x 10^10^ Virus-Like-Particles/mL (FigureS1). The mice were subjected to an oral glucose tolerance test (OGTT)[18] at week 12 of the study and food intake and mouse weight was monitored. The study was approved by the Danish Competent Authority, The Animal Experimentation Inspectorate, under the Ministry of Environment and Food of Denmark, and performed under license No. 2017-15-0201-01262 C1-3. Procedures were carried out in accordance with the Directive 2010/63/EU and the Danish law LBK Nr 726 af 09/091993, and housing conditions as earlier described[17]. Blood plasma, intestinal content from the cecum and colon as well as tissue from the liver and ileum were sampled at termination at week 13 (18 weeks old). Mouse data (weight, OGTT levels, etc.) were analysed in GraphPad Prism using one-way ANOVA with Tukeys test.

### Pre-processing of faecal samples

This study included in total 79 intestinal content samples, of which 40 were isolated from cecum and 39 from colon. Pre-processing was performed as earlier described[17].

### Bacterial DNA extraction, sequencing, and pre-processing of raw data

Tag-encoded 16S rRNA gene amplicon sequencing was performed on an Illumina NextSeq using v2 MID output 2×150 cycles chemistry (Illumina, CA, USA). DNA extraction and library building for amplicon sequencing was performed in accordance with Krych *et al*.[19]. The average sequencing depth (Accession: PRJEB32560, available at ENA) for the cecum 16S rRNA gene amplicons was 164,147 reads (min. 22,732 reads and max. 200,2003 reads) and 166,012 reads for colon (min. 89,528 reads and max. 207,924 reads) (TableS1). bOTU tables (bacterial-operational taxonomic units) were generated and taxonomy assigned as earlier described[17]. Bacterial density of the caecal and colon content was estimated by quantitative real-time polymerase chain reaction (qPCR) as previously described[20], using the 16S rRNA gene primers (V3 region) also applied for the amplicon sequencing[19].

### Viral DNA extraction, sequencing and pre-processing of raw data

The enteric viral community was purified, DNA extracted and the gut metavirome determined as previously described[17]. The average sequencing depth (Accession: PRJEB32560, available at ENA) for the cecum viral metagenome was 612,640 reads/sample (min. 277,582 reads and max. 1,219,178 reads) and 356,976 reads/sample for colon (min. 33,773 reads and max. 584,681 reads) (TableS1). For each sample, reads were treated with Trimmomatic[21] and Usearch[22] and subjected to within-sample *de novo* assembly with MetaSpades v.3.5.0[23,24]. Viral contigs were identified with Kraken2[25], VirFinder[26], PHASTER[27], and virus orthologous proteins (www.vogdb.org). Contaminations of non-viral contigs like bacteria, human, mice, and plant DNA were removed (FigureS2). The remaining contigs constituted the vOTU (viral-operational taxonomic unit) table.

### Gene expression assay

Genes investigated for expression analysis were selected based on relevant pathways for each tissue. For the liver, genes involved in metabolic pathways (triglyceride, carbohydrate, bile and cholesterol metabolism) and inflammation were selected. For the ileum, genes involved in inflammation, gut microbiota signalling and gut barrier function were selected [28,29]. Primer sequences are listed in Table S2. qPCR was performed using the Biomark HD system (Fluidigm Corporation) on 2x 96.96 IFC chips on pre-amplified cDNA duplicates following the instructions of the manufacturer with minor adjustments as previously described[30]. Gene expression data was analysed in R using linear models (one-way ANOVA) with either HF or LF as control groups.

### Blood plasma metabolome analysis

Plasma samples were prepared for ultra-performance liquid chromatography mass spectrometry (UPLC-MS) analysis according to a previously published protocol[31]. Principal component analysis (PCA) and orthogonal projection to latent structures-discriminant analysis (OPLS-DA) models were built on the plasma metabolic profiles to identify biochemical variation between the groups. Among the features driving the different OPLS-DA models, only those with variable importance in the projection (VIP) scores > 2 were further investigated. Putative annotation was achieved through searching for the m/z values in online databases such as HMDB (http://www.hmdb.ca), METLIN (http://metlin.scripps.edu) and Lipidmaps (http://www.lipidmaps.org). Additionally, fragmentation patterns derived from MS^e^ experiment were compared to online spectra when available.

### Bioinformatic analysis of bacterial and viral DNA

Prior to any analysis the raw read counts in vOTU-tables were normalised by reads per kilo base per million mapped reads (RPKM)[32]. b- and vOTU’s that were detected in less than 8% of the samples were discarded to reduce noise while still maintaining an average total abundance close to 98%. ANOSIM and Kruskal Wallis was used to evaluate multiple group comparisons. Regularised Canonical Correlation Analysis (rCCA) was performed with mixOmics v. 6.8.0. R-package[33] to predict correlations between bacterial and viral taxa. Only vOTUs ≥ 5000 bp were included in the rCCA. The machine learning algorithm random forest[34] was applied to select variables explaining the dataset and normalised in range of -1:1 ((x-mean)/max(abs(x-mean)) and visualised by Heatmap3[35].

## RESULTS

Here we investigated the potential of faecal virome transplantation (FVT) to shift the phenotype of obesity and T2D in DIO male C57BL/6NTac mice towards a lean mice phenotype. Intestinal contents from the cecum and colon were isolated, but here only results from the cecum samples are be reported. Complete equivalent analysis of colon samples can be found in Figure S10.

### FVT from lean donor decreases weight gain and normalises blood glucose tolerance in DIO mice

Mice were weighed pre and post FVT with 1-2 weeks of interval. At both 4- and 6-weeks post first FVT (15 and 17 weeks old) a significantly lower body weight gain was observed in the HF+FVT (p < 0.017) and the HF+Amp mice (p < 0.006) compared to the HF mice (Figure 2a). Intriguingly, OGTT showed no significant differences (p > 0.842) between the LF and the HF+FVT mice, while the OGTT level of the HF mice was significantly increased (p < 0.0001) compared to both the LF and HF+FVT group suggesting that FVT had normalised the blood glucose tolerance in the HF+FVT mice (Figure 2b). Furthermore, the OGTT of HF+Amp+FVT was comparable to the HF mice (p > 0.999), indicating that the initial disruption of the bacterial composition by the ampicillin treatment had counteracted the effect of the FVT in the HF+Amp+FVT mice. Which simultaneously suggest that the effects associated to the FVT occurs via alterations in the GM component. Non-fasted blood glucose was measured regularly in addition to HbA1c levels and food consumption pr. cage (FigureS4).

**Figure 2:**
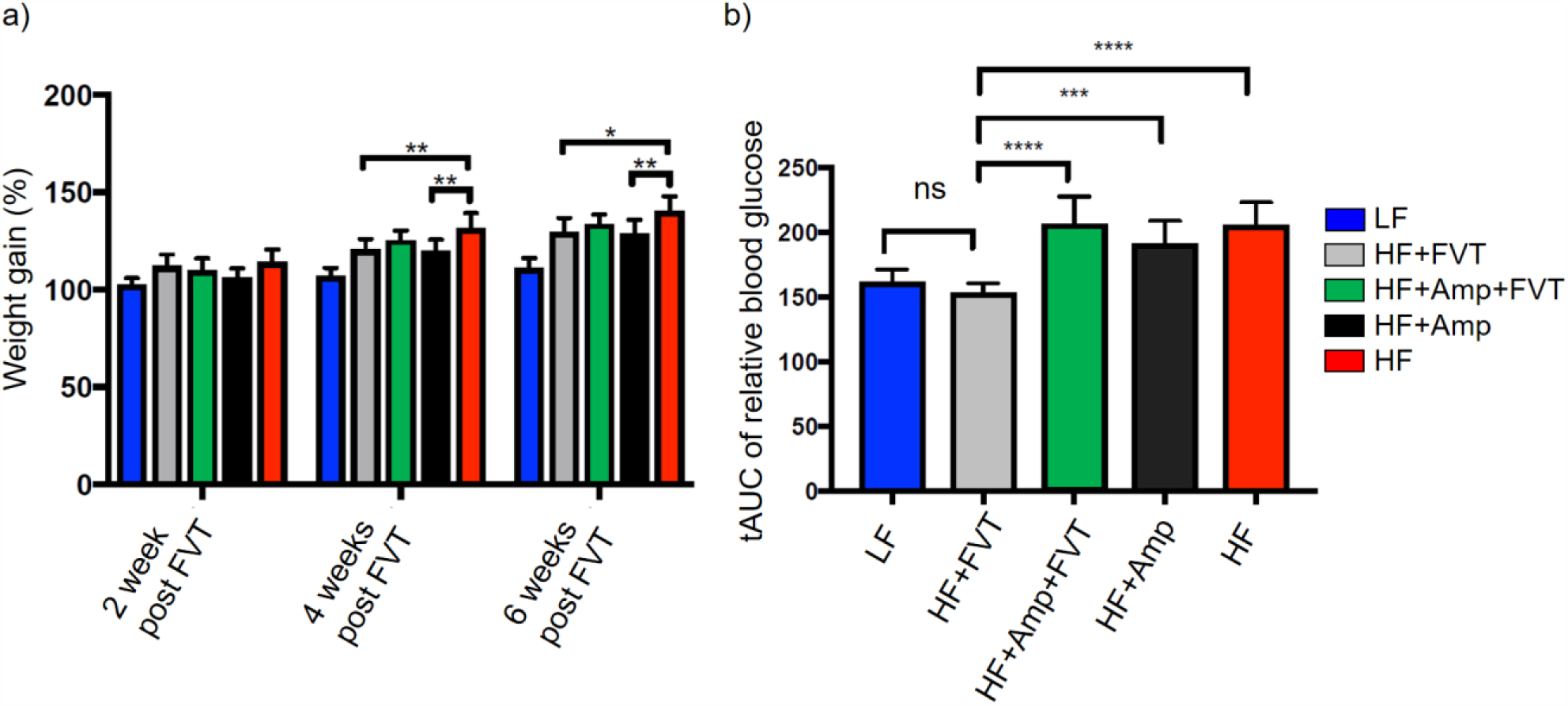
a) Bar plot of body weight gain measured at 2, 4, and 6 weeks (13, 15, 17 weeks old respectively) post first faecal virome transplantation (FVT). b) Oral glucose tolerance test (OGTT) levels measured 6 weeks post first FVT (17 weeks old). Values are based on the total area under curve (tAUC) relatively to the blood glucose levels of the individual mouse. Significant differences of the pairwise comparison at week 4 and 6 post first FVT was excluded in the figures to increase the visualisation. HF = high-fat, LF = low-fat, Amp = ampicillin, ns = not significant. * = p < 0.05, ** = p < 0.006, *** = p < 0.0005, **** = p < 0.0001.

### FVT enhances the expression of genes involved in whole-body and energy homeostasis

Gene expression panels were designed to target genes relevant for obesity and T2D in liver and ileum tissue to measure if the expression of these genes in the HF+FVT mice were significantly different to the HF mice, while being comparable to the LF mice. These conditions were represented by several genes involved in whole-body and energy homeostasis (Figure 3). Gene expression analysis suggested that the FVT had blunted the differences in gene expression caused by the HF diet resulting in an expression profile closer resembling that of the healthy LF mice. The expression levels of *Lepr*^*Liver*^, *Ffar2*^*Ileum*^, *Klb*^*Liver*^, *Ppargc1a*^*Liver*^, and *Igfbp2* ^*Liver*^ were significantly increased in the HF+FVT compared to the HF mice, whereas *Socs3*^*Liver*^ and *Myc*^*Liver*^ were significantly decreased (Figure 3). Generally, the gene expression levels (except *Socs3*^*Liver*^) in the HF+FVT mice fell between the levels observed in the LF and HF mice. Significant differences (p < 0.05) were observed in the expression of several other genes (57 of 74 for liver tissue and 58 of 74 for ileum tissue) between the experimental groups (see TableS3 for complete list).

**Figure 3:**
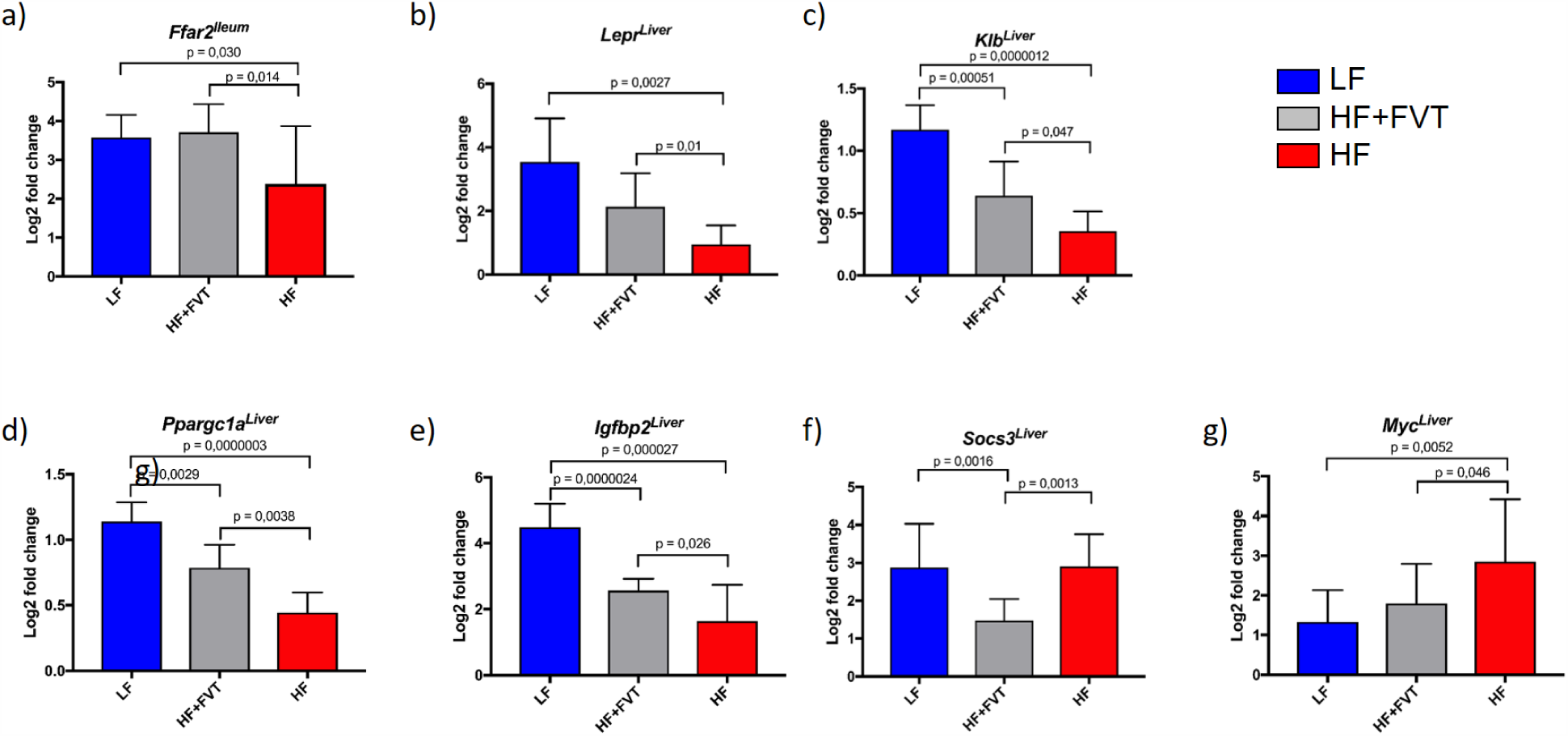
Gene expression levels at termination (18 weeks old) of a) Lepr^Liver^ (leptin cytokine receptors), b) Ffar2^Ileum^ (free fatty acid receptor), c) Klb^Liver^ (Beta-klotho), d) Ppargc1a^Liver^ (Peroxisome proliferator-activated receptor gamma coactivator 1-alpha), e) Igfbp2 ^Liver^(insulin like growth factor binding protein), f) Socs3^Liver^ (suppressor of cytokine signalling), and g) Myc^Liver^ (transcription factor). Linear models (one-way ANOVA) with either HF or LF as control groups was performed to calculate group significance. The log2 fold change is a measure of the relative gene expression and is based on log2 transformed expression values normalised to the sample with the lowest value. HF = high-fat, LF = low-fat, Amp = ampicillin, FVT = faecal virome transplantation.

### FVT-mediated shift in the gut microbiota component

At termination the number of 16S rRNA gene copies/g of the cecum samples varied from 1.46 x 10^10^ – 2.70 × 10^10^ (FigureS3). The bacterial Shannon diversity index of the LF mice was significantly higher than the HF mice (p < 0.005) but similar to the HF+FVT mice (p = 0.816). The ampicillin-treated HF+Amp mice had the lowest Shannon diversity index of all groups at termination (i.e. 7 weeks after treatment with ampicillin) – and FVT increased the Shannon diversity index of ampicillin treated HF+Amp+FVT mice (p < 0.016) (Figure 4a). The FVT had no significant (p > 0.09) impact on the viral Shannon index, whereas the ampicillin treatment significantly (p < 0.0003) increased the viral Shannon index (Figure 4b), possibly due to induction of prophages (FigureS6).

**Figure 4:**
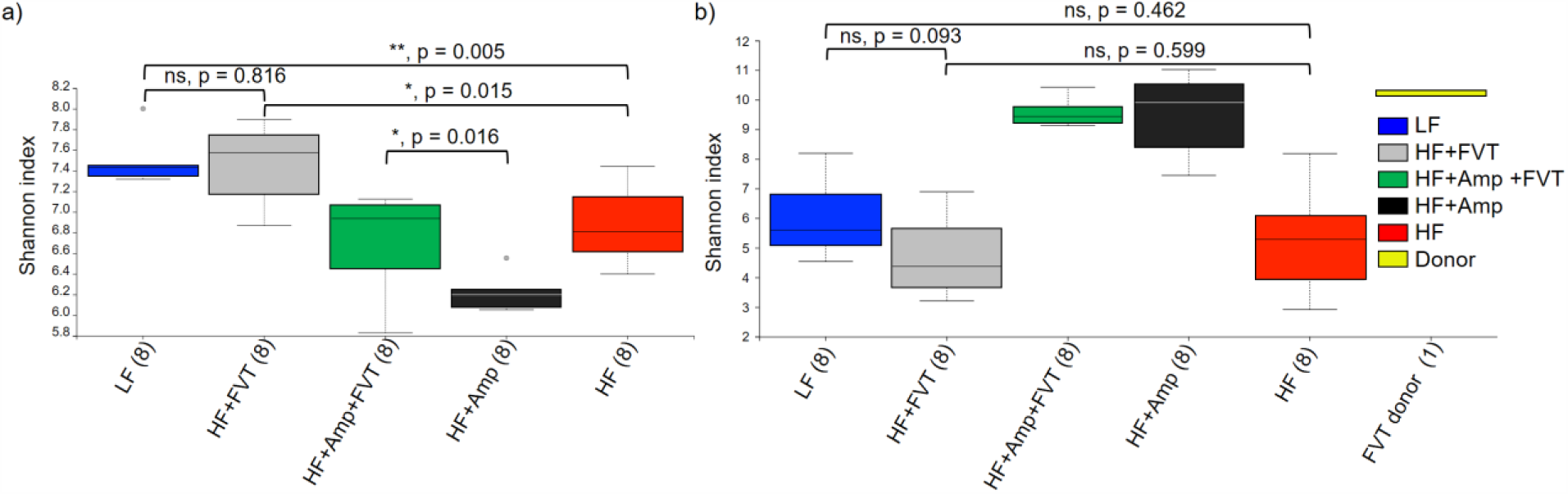
Shannon index of the caecal a) bacterial and b) viral community at termination (18 weeks old). The parentheses show the number of samples from each group included in the plot and grey dots indicate outliers. HF = high-fat, LF = low-fat, Amp = ampicillin, FVT = faecal virome transplantation, ns = not significant. * = p < 0.05, ** = p < 0.0051.

The FVT strongly influenced both the bacterial (Figure 5a, p < 0.003) and viral (Figure 5b, p < 0.001) composition as determined by the Bray-Curtis dissimilarity-metric, as illustrated by a clear separation of the HF+FVT compared with the HF mice and HF+Amp+FVT with the HF+Amp mice. Further, all experimental groups were pairwise significantly separated (p < 0.003) in both the viral and bacterial community (TableS4), including LF vs. HF+FVT (p < 0.001). Overall, our findings suggest that FVT strongly influenced and partly reshape the GM composition both with and without ampicillin treatment.

**Figure 5:**
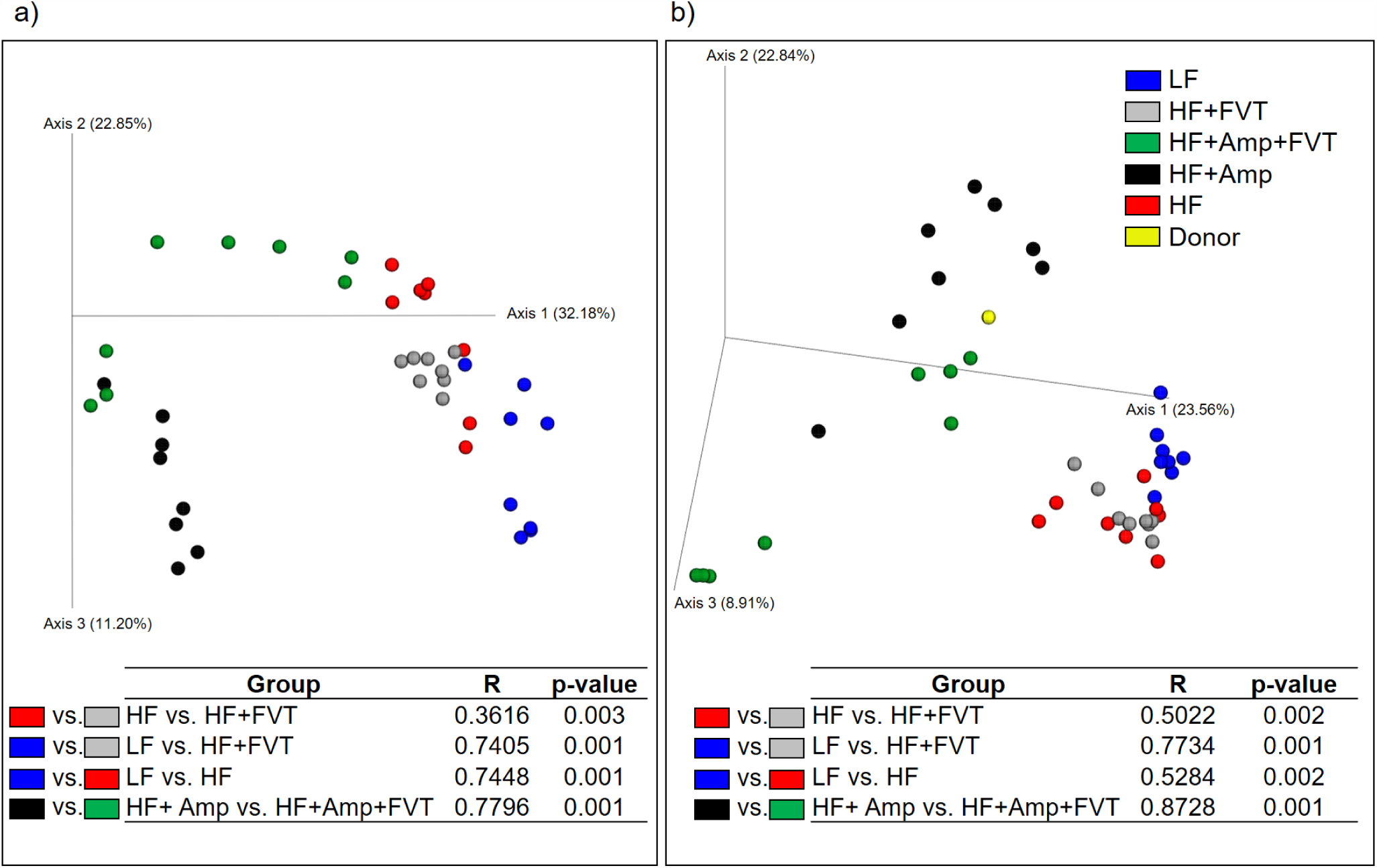
Bray Curtis dissimilarity metric PCoA based plots of a) the caecal bacterial community and b) viral community at termination (18 weeks old). Tables include ANOSIM of the Bray Curtis dissimilarity to show the effect of the faecal virome transplantation (FVT) on the gut microbiota (GM) composition. HF = high-fat, LF = low-fat, Amp = ampicillin.

### FVT-mediated shift in the blood plasma metabolome profile

The influence of FVT on host metabolome was determined by untargeted UPLC-MS analysis of plasma samples. A PCA model was built on a refined dataset comparing LF, HF and HF+FVT profiles (Figure 6 and FigureS5 for all groups). Consistent with the other measures, the plasma profiles of HF+FVT mice were positioned between the HF and LF mice. Pairwise OPLS-DA models were constructed and all the models (LF vs. HF, LF vs. HF+FVT, HF vs. HF+FVT) were significant (p < 0.025), supporting the separation of the three groups. Among the selected features with a VIP score > 2, only those correlating with relevant gene expression (based on rCCA), bacterial or viral abundance were further investigated for annotation. These features investigated consisted mainly of saturated and unsaturated lyso-phosphatidylcholine (LysoPC) and/or phosphatidylcholines (PCs), whereas the remaining features consisted of varieties of amino acids or unidentifiable metabolites (Table S5). Overall the HF mice had higher levels of LysoPC(18:2), LysoPC(22:2), and PC(16:0/22:6) than the LF mice which on the other hand had higher plasma levels of LysoPC(22:4) and PC(18:1/O-18:2). The HF+FVT mice had increased levels of circulating LysoPC(16:0), LysoPC(18:2), and PC(16:0/22:6) compared to the LF mice while the levels of LysoPC(22:4) and PC(18:1/O-18:2) were decreased. Further, the HF+FVT mice appeared with higher levels of LysoPC(16:0), LysoPC(18:0), and PC(18:1/O-18:2) compared to the HF mice, whereas the LF mice showed higher levels of LysoPC(22:4) and PC(18:1/O-18:2) compared to the HF mice.

**Figure 6:**
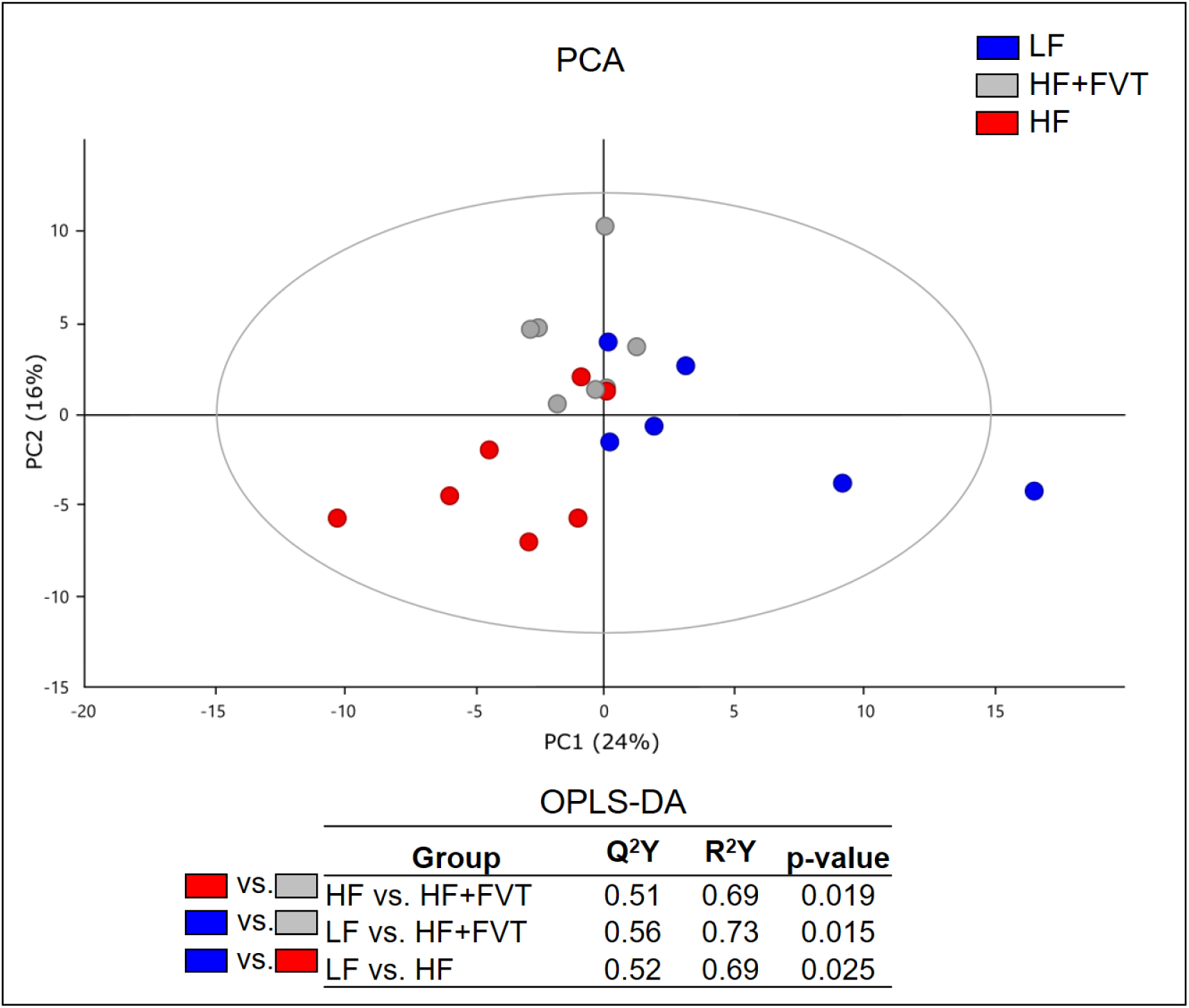
PCA scores plot obtained from ESI+ UPLC-MS of plasma from LF, HF and HF+FVT (R2=0.40 and Q2=0.11) at termination (18 weeks old). Table includes supervised OPLS-DA models generated by pairwise comparisons. HF = high-fat, LF = low-fat, Amp = ampicillin.

## DISCUSSION

FVT has been successfully used to treat rCDI[14], an effect probably mediated by the ability of FVT to reshape the GM of the recipient. However, several parameters differentiate obesity and T2D from rCDI to which is characterised by a clonal infection highly susceptible to viral attack, a highly dysbiotic GM and the extensive use of antibiotics[14]. Here, we show that FVT from lean mice donors to obese recipients successfully counteracts some of the adverse effects of a HF diet with a decrease in body weight gain (p < 0.0034) and a normalised blood glucose tolerance relative to mice fed a LF diet. Indeed, HF+FVT mice showed exactly the same response in an OGTT as the LF mice (p > 0.842, Figure 2), whereas similar effects were not observed in the HF+Amp+FVT mice. As expected[36], the diversity of the caecal bacterial community (α-diversity) in the HF mice was significantly (p < 0.005) lower compared to LF mice, but surprisingly the bacterial diversity of the HF+FVT was similar to the LF (p > 0.81) and elevated in comparison (p < 0.015) to the HF mice. No significant viral diversity differences were observed between LF vs. HF, LF vs. HF+FVT, and HF vs. HF+FVT. The significantly increased (p < 0.003) diversity of the viral community in the ampicillin treated mice (Figure 4b) was likely due to induction of prophages[37], as indicated by an increased (p < 0.05) presence of viral contigs containing integrase genes in the ampicillin treated groups (FigureS6). The composition of the viral community was dominated by order Caudovirales and family *Microviridae* viruses, which is in accordance with former studies investigating the enteric viral community in mammals[9,12]. The FVT strongly influenced both the bacterial and viral GM composition (Figure 5), with HF+FVT being significantly (p < 0.003) different from both the HF as well as the LF mice. The highly diverse donor virome[17] as well as the distinct nutrition profiles (HF-diet vs. LF-diet) most likely explain the divergence in the GM composition profiles between the HF+FVT and the LF mice.

Gene expression analysis of obesity and T2D associated genes in liver and ileum showed that seven relevant genes (Figure 3) were differentially expressed (p < 0.05) between HF+FVT and HF mice. The *Ffar2* gene is involved in energy homeostasis and can influence glucose homeostasis through GLP-1 regulation and leptin production[38,39], which might lower food intake and via GLP-1 decrease blood glucose levels by enhancing the production of insulin[38]. The *Ffar2* gene may also influence whole-body homeostasis through regulation of adipogenesis and lipid storage of adipocytes[40]. The expression of *Ffar2*^*Ileum*^ was comparable between LF and HF+FVT while in the HF mice expression was clearly reduced (Figure 3a). The *Lepr* genes are involved in the regulation of food intake, and *Lepr* deficient subjects rapidly increase body weight[41]. *Klb* contributes to the repression of cholesterol 7-alpha-hydroxylase and thereby regulate bile acid synthesis[42] with reduced expression in obese subjects[43]. *Ppargc1a* expression has been reported to be reduced and linked with islet insulin secretion in T2D patients[44], as well as playing a pivotal role in regulating energy homeostasis[45]. *Igfbp2* is involved in the insulin-like growth factor-axis influencing cell growth and proliferation[46] and high expression levels have been linked to the protection against T2D in human studies[47]. The gene expression of *Lepr*^*Liver*^ (Figure3b), *Klb*^*Liver*^ (Figure3c), *Ppargc1a*^*Liver*^ (Figure3d), and *Igfbp2*^*Liver*^ (Figure3e) were all significantly increased in the LF and HF+FVT mice compared to the HF mice. Knock-out of *Socs* genes have been reported to prevent insulin resistance in obesity as the result of a decrease in ceramide synthesis[48]. The *Socs3*^*Liver*^ levels were significantly (p < 0.0001) decreased in HF+FVT compared to both HF and LF mice (Figure 3f). However, the overall *Socs3*^*Liver*^ levels in the LF mice was affected by high inter-group variation. The expression of *Myc* has been found to be increased in mice provided HF diet, and a decrease in bodyweight was obtained with haploin-sufficient mice (*c-Myc*^*+/-*^)[49]. The expression of *Myc*^*Liver*^ was comparable between LF and HF+FVT while in the HF mice expression was clearly increased (Figure 3g). Overall, these findings indicate that FVT treatment affects the expression of genes involved in stimulating appetite, blood glucose tolerance, and whole-body and energy homeostasis.

An extensive study of metabolic syndrome in humans, showed strong correlations between certain blood plasma metabolites and the GM component[50], and some of these correlations appear specific to the pre-diabetic state[50]. Furthermore, blood plasma metabolome seem to predict the GM diversity[51]. The blood plasma metabolome profile of the HF+FVT mice differed significantly (p < 0.025) from both the LF and HF mice. SCFAs are expected to regulate the *Ffar2* gene[52] but our feature annotation did not detect any clear differences in circulating SCFAs between treatment groups. Further, LysoPCs are reduced in obesity and T2D[53], and we indeed observed a decrease in LysoPCs levels when comparing the HF with HF+FVT group and HF with the LF group, although the results were not clear. The beneficial effects associated to FVT were knocked down by the ampicillin treatment (except for weight gain), which was also reduced in the HF+Amp relative to HF, but the FVT consistently reshaped the GM composition profile in the HF+Amp+FVT mice as well.

Regularized Canonical Correlation Analysis (rCCA) suggested potential host-phage pair relations by strong (r > 0.75) positive and negative correlations between certain bacterial (order Bacteroidales, Clostridiales) and viral (order Caudovirales, family *Microviridae* and uncharacterised viruses) taxa. Likewise, random forest selected variables showed bacterial and viral co-abundance profiles that differentiated the five experimental groups (Figure 7) supporting the effect of the FVT on the GM component.

**Figure 7:**
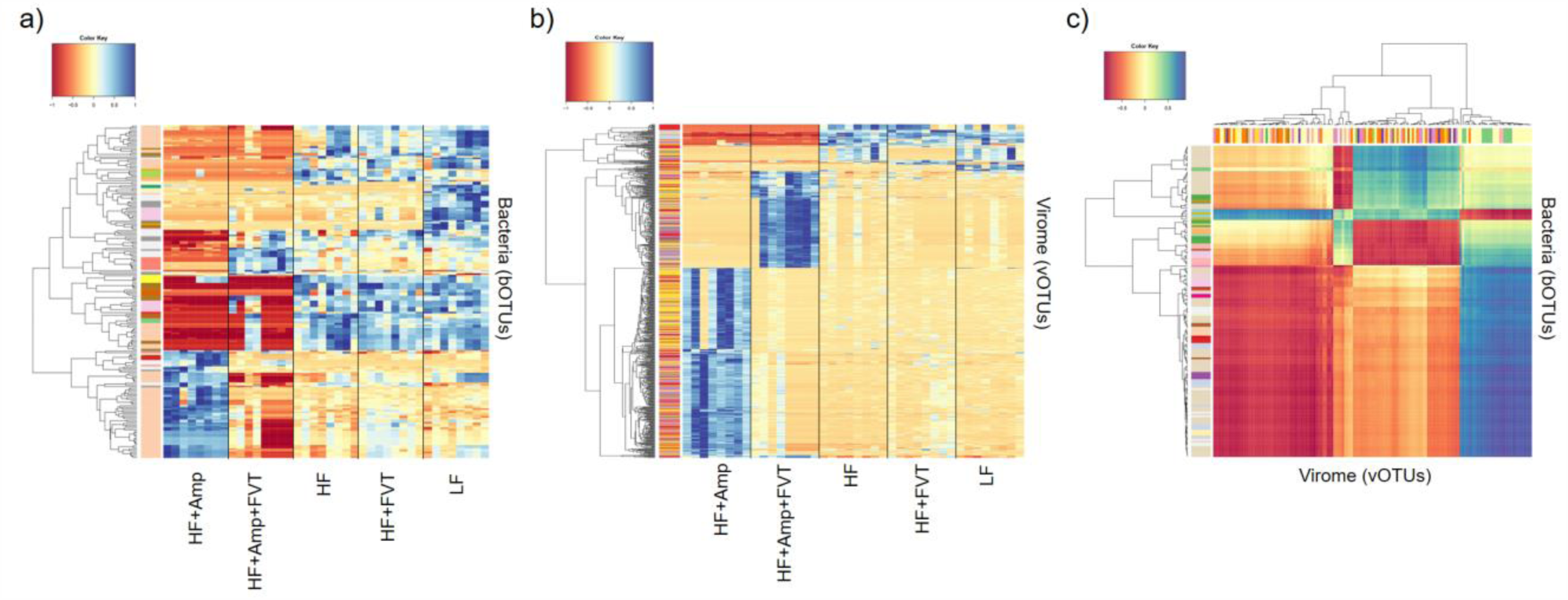
Heatmaps illustrating the bacterial a) and viral b) profile of all five experimental groups, as well as strong correlations between certain clusters of bacteria and viruses c). Detailed information and high-resolution images can be found as supplemental materials in Figure S7 and S8.

Zuo *et al.*[54] investigated how the phage community could be related to the effect of FMT against rCDI and found that a few patients did not respond on the treatment. The common denominator of the patients that did respond on the FMT treatment was that the Caudovirales richness in the donor faeces was higher than the Caudovirales richness in the recipient, whereas most of the non-responder recipients had a higher Caudovirales richness than the donor. Based on these findings, Zuo *et al.* hypothesised that a higher Caudovirales richness in the donor compared to the recipient is important for a successful FMT treatment[54]. The Caudovirales richness of the FVT donor virome in our study (faecal virome from mice purchased at three different vendors[17]) was higher than any of the recipient groups at termination (FigureS9). Whereas the Caudovirales richness of the viromes from the individual mouse vendors were notably less compared to the FVT virome and both recipient groups. Although further studies are needed, these findings are consistent with those of Zuo *et al.* and suggests that a virome from a single vendor/donor might not have the same effect as when multiple viromes are combined for FVT.

We expect that phages transferred with the FVT in the HF+FVT mice have reshaped the GM component and thereby shifted the phenotype of the obese HF mice to be in closer resemblance of the lean LF mice. This hypothesis seems plausible since several recent studies have reported a correlation between phage diversity and intestinal microbiome diversity[12], a FVT-mediated restoration of the GM of antibiotic treated mice[15], and a phage-mediated shift in the gut metabolome profile[13].

In conclusion, we here demonstrate the use of FVT targeting obesity and T2D in an animal model. Although the study is a proof-of-concept, our findings highlight the potential of using phage-mediated therapy against obesity and T2D[6] that represents a world-wide health threat[1].

## Acknowledgement

Thanks to Julie Mou Larsen for assisting with sampling of mouse tissue and associated method description. In addition, we thank Helene Farlov, Mette Nelander at Section of Experimental Animal Models and Liv de Vries at Section for Microbiology and Fermentation (University of Copenhagen, Denmark) for taking care of the animals. We thank Helle Keinicke and Marina Kjærgaard Gerstenberg for selecting the genes and providing the primers for the liver gene expression panel.

## Contributors

T.S.R., C.M.M., A.K.H., D.S.N, and F.K.V. conceived the research idea and designed the study; C.M.M. and T.S.R. performed the animal experiments; T.S.R., C.M.M., and S.Z. processed the samples in the laboratory; T.S.R., C.M.M., L.H.H., W.K., D.S.N, J.S., S.Z., J.C.M., and F.K.V performed data analysis; T.S.R. and D.S.N was responsible for the first and final draft of the manuscript, as well as all requested revisions. All authors critically revised and approved the final version of the manuscript.

## Competing interests

All authors declare no conflicts of interest.

## Funding

Funding was provided by the Danish Council for Independent Research with grant ID: DFF-6111-00316 (PhageGut) and the Danish Innovation Fund project - 7076-00129B, MICROHEALTH.

## References

1 Leitner DR, Frühbeck G, Yumuk V, et al. Obesity and type 2 diabetes: Two diseases with a need for combined treatment strategies - EASO can lead the way. Obes Facts 2017;10:483–92. doi:10.1159/000480525

2 Maruvada P, Leone V, Kaplan LM, et al. The Human Microbiome and Obesity: Moving beyond Associations. Cell Host Microbe 2017;22:589–99. doi:10.1016/j.chom.2017.10.005

3 Cho I, Yamanishi S, Cox L, et al. Antibiotics in early life alter the murine colonic microbiome and adiposity. Nature 2012;488:621–6. doi:10.1038/nature11400

4 Ridaura VK, Faith JJ, Rey FE, et al. Gut microbiota from twins discordant for obesity modulate metabolism in mice. Science (80-) 2013;341:1241214. doi:10.1126/science.1241214

5 Kelly CR, Kahn S, Kashyap P, et al. Update on Fecal Microbiota Transplantation 2015: Indications, Methodologies, Mechanisms, and Outlook. Gastroenterology 2015;149:223–37. doi:10.1053/j.gastro.2015.05.008

6 Gupta S, Allen-vercoe E, Petrof EO. Fecal microbiota transplantation?: in perspective. Therap Adv Gastroenterol 2016;9:229–39. doi:10.1177/1756283X15607414

7 Wang JYJWJYJW, Kuo CH, Kuo FC, et al. Fecal microbiota transplantation: Review and update. J Formos Med Assoc 2019;118:S23–31. doi:10.1016/j.jfma.2018.08.011

8 FDA. FDA - Important Safety Alert Regarding Use of Fecal Microbiota for Transplantation and Risk of Serious Adverse Reactions Due to Transmission of Multi-Drug Resistant Organisms. 2019.https://www.fda.gov/vaccines-blood-biologics/safety-availability-biologics/important-safety-alert-regarding-use-fecal-microbiota-transplantation-and-risk-serious-adverse (accessed 4 Jul 2019).

9 Reyes A, Haynes M, Hanson N, et al. Viruses in the faecal microbiota of monozygotic twins and their mothers. Nature 2011;466:334–8. doi:10.1038/nature09199

10 Ross A, Ward S, Hyman P. More Is Better?: Selecting for Broad Host Range Bacteriophages. 2016;7:1–6. doi:10.3389/fmicb.2016.01352

11 Howe A, Ringus DL, Williams RJ, et al. Divergent responses of viral and bacterial communities in the gut microbiome to dietary disturbances in mice. ISME J 2016;10:1217–27. doi:10.1038/ismej.2015.183

12 Moreno-Gallego JL, Chou SP, Di Rienzi SC, et al. Virome Diversity Correlates with Intestinal Microbiome Diversity in Adult Monozygotic Twins. Cell Host Microbe 2019;25:261-272.e5. doi:10.1016/j.chom.2019.01.019

13 Hsu BB, Gibson TE, Yeliseyev V, et al. Dynamic Modulation of the Gut Microbiota and Metabolome by Bacteriophages in a Mouse Model. Cell Host Microbe 2019;25:803-814.e5. doi:10.1016/j.chom.2019.05.001

14 Ott SJ, Waetzig GH, Rehman A, et al. Efficacy of Sterile Fecal Filtrate Transfer for Treating Patients With Clostridium difficile Infection. Gastroenterology 2017;152:799–811. doi:10.1053/j.gastro.2016.11.010

15 Draper LA, Ryan FJ, Dalmasso M, et al. Autochthonous faecal virome transplantation (FVT) reshapes the murine microbiome after antibiotic perturbation. bioRxiv 2019;:591099. doi:10.1101/591099

16 Fraulob JC, Ogg-Diamantino R, Fernandes-Santos C, et al. A Mouse Model of Metabolic Syndrome: Insulin Resistance, Fatty Liver and Non-Alcoholic Fatty Pancreas Disease (NAFPD) in C57BL/6 Mice Fed a High Fat Diet. J Clin Biochem Nutr 2010;46:212–23. doi:10.3164/jcbn.09-83

17 Rasmussen TS, de Vries L, Kot W, et al. Mouse Vendor Influence on the Bacterial and Viral Gut Composition Exceeds the Effect of Diet. Viruses 2019;11:435. doi:10.3390/v11050435

18 Rune I, Rolin B, Lykkesfeldt J, et al. Long-term Western diet fed apolipoprotein E-deficient rats exhibit only modest early atherosclerotic characteristics. Sci Rep 2018;8:1–12. doi:10.1038/s41598-018-23835-z

19 Krych L, Kot W, Bendtsen KMB, et al. Have you tried spermine? A rapid and cost-e ff ective method to eliminate dextran sodium sulfate inhibition of PCR and RT-PCR. J Microbiol Methods J 2018;144:1–7. doi:10.1016/j.mimet.2017.10.015

20 Ellekilde M, Krych L, Hansen CHFHF, et al. Characterization of the gut microbiota in leptin deficient obese mice - Correlation to inflammatory and diabetic parameters. Res Vet Sci 2014;96:241–50. doi:10.1016/j.rvsc.2014.01.007

21 Bolger AM, Lohse M, Usadel B. Trimmomatic: A flexible trimmer for Illumina sequence data. Bioinformatics 2014;30:2114–20. doi:10.1093/bioinformatics/btu170

22 Edgar RC. UPARSE: highly accurate OTU sequences from microbial amplicon reads. Nat Methods 2013;10:996–8. doi:10.1038/nmeth.2604

23 Bankevich A, Nurk S, Antipov D, et al. SPAdes: A new genome assembly algorithm and its applications to single-cell sequencing. J Comput Biol 2012;19:455–77. doi:10.1089/cmb.2012.0021

24 Nurk S, Meleshko D, Korobeynikov A, et al. metaSPAdes: a new versatile metagenomic assembler. Genome Res 2017;27:824–34. doi:10.1101/gr.213959.116

25 Wood DE, Salzberg SL. Kraken: ultrafast metagenomic sequence classification using exact alignments. Genome Biol 2014;15:R46. doi:10.1186/gb-2014-15-3-r46

26 Ren J, Ahlgren NA, Lu YY, et al. VirFinder: a novel k-mer based tool for identifying viral sequences from assembled metagenomic data. Microbiome 2017;5:69. doi:10.1186/s40168-017-0283-5

27 Arndt D, Grant JR, Marcu A, et al. PHASTER: a better, faster version of the PHAST phage search tool. Nucleic Acids Res 2016;44:1–6. doi:10.1093/nar/gkw387

28 Bendtsen KM, Hansen CHF, Krych Ł, et al. Immunological effects of reduced mucosal integrity in the early life of BALB/c mice. PLoS One 2017;12:1–20. doi:10.1371/journal.pone.0176662

29 Zachariassen LF, Krych L, Rasmussen SH, et al. Cesarean Section Induces Microbiota-Regulated Immune Disturbances in C57BL/6 Mice. J Immunol 2019;202:142–50. doi:10.4049/jimmunol.1800666

30 Mentzel CMJ, Cardoso TF, Pipper CB, et al. Deregulation of obesity-relevant genes is associated with progression in BMI and the amount of adipose tissue in pigs. Mol Genet Genomics 2018;293:129–36. doi:10.1007/s00438-017-1369-2

31 Sarafian MH, Gaudin M, Lewis MR, et al. Objective Set of Criteria for Optimization of Sample Preparation Procedures for Ultra-High Throughput Untargeted Blood Plasma Lipid Profiling by Ultra Performance Liquid Chromatography-Mass Spectrometry. Anal Chem 2014;86:5766–74. doi:10.1021/ac500317c

32 Roux S, Emerson JB, Eloe-Fadrosh EA, et al. Benchmarking viromics: an *in silico* evaluation of metagenome-enabled estimates of viral community composition and diversity. PeerJ 2017;5:e3817. doi:10.7717/peerj.3817

33 Rohart F, Gautier B, Singh A, et al. mixOmics: An R package for ‘omics feature selection and multiple data integration. PLOS Comput Biol 2017;13:e1005752. doi:10.1371/journal.pcbi.1005752

34 Breiman L. Random forests. Mach Learn 2001;45:5–32. doi:10.1023/A:1010933404324

35 Zhao S, Guo Y, Sheng Q, et al. Heatmap3: An improved heatmap package with more powerful and convenient features. BMC Bioinformatics 2014;15:P16. doi:10.1186/1471-2105-15-S10-P16

36 Sun L, Ma L, Ma Y, et al. Insights into the role of gut microbiota in obesity: pathogenesis, mechanisms, and therapeutic perspectives. Protein Cell 2018;9:397–403. doi:10.1007/s13238-018-0546-3

37 Allen HK, Looft T, Bayles DO, et al. Antibiotics in feed induce prophages in swine fecal microbiomes. MBio 2011;2:1–9. doi:10.1128/mBio.00260-11

38 Hudson BD, Due-Hansen ME, Christiansen E, et al. Defining the molecular basis for the first potent and selective orthosteric agonists of the FFA2 free fatty acid receptor. J Biol Chem 2013;288:17296–312. doi:10.1074/jbc.M113.455337

39 Ichimura A, Hasegawa S, Kasubuchi M, et al. Free fatty acid receptors as therapeutic targets for the treatment of diabetes. Front Pharmacol 2014;5:1–6. doi:10.3389/fphar.2014.00236

40 Murdock PR, Pike NB, Eilert MM, et al. The Orphan G Protein-coupled Receptors GPR41 and GPR43 Are Activated by Propionate and Other Short Chain Carboxylic Acids. J Biol Chem 2003;278:11312–9. doi:10.1074/jbc.m211609200

41 Dubern B, Clement K. Leptin and leptin receptor-related monogenic obesity. Biochimie 2012;94:2111–5. doi:10.1016/j.biochi.2012.05.010

42 Ito S, Fujimori T, Furuya A, et al. Impaired negative feedback suppression of bile acid synthesis in mice lacking βKlotho. J Clin Invest 2005;115:2202–8. doi:10.1172/JCI23076

43 Kruse R, Vienberg SG, Vind BF, et al. Effects of insulin and exercise training on FGF21, its receptors and target genes in obesity and type 2 diabetes. Diabetologia 2017;60:2042–51. doi:10.1007/s00125-017-4373-5

44 Ling C, Del Guerra S, Lupi R, et al. Epigenetic regulation of PPARGC1A in human type 2 diabetic islets and effect on insulin secretion. Diabetologia 2008;51:615–22. doi:10.1007/s00125-007-0916-5

45 Charos AE, Reed BD, Raha D, et al. A highly integrated and complex PPARGC1A transcription factor binding network in HepG2 cells. Genome Res 2012;22:1668–79. doi:10.1101/gr.127761.111

46 Rajpathak SN, Gunter MJ, Wylie-Rosett J, et al. The role of insulin-like growth factor-I and its binding proteins in glucose homeostasis and type 2 diabetes. Diabetes Metab Res Rev 2009;25:3–12. doi:10.1002/dmrr.919

47 Wittenbecher C, Ouni M, Kuxhaus O, et al. Insulin-like growth factor binding protein (IGFBP-2) and the risk of developing type 2 diabetes. Diabetes 2019;68:188–97. doi:10.2337/db18-0620

48 Yang G, Badeanlou L, Bielawski J, et al. Central role of ceramide biosynthesis in body weight regulation, energy metabolism, and the metabolic syndrome. Am J Physiol Metab 2009;297:E211–24. doi:10.1152/ajpendo.91014.2008

49 Liu S, Kim TH, Franklin DA, et al. Protection against High-Fat-Diet-Induced Obesity in MDM2C305F Mice Due to Reduced p53 Activity and Enhanced Energy Expenditure. Cell Rep 2017;18:1005–18. doi:10.1016/j.celrep.2016.12.086

50 Org E, Blum Y, Kasela S, et al. Relationships between gut microbiota, plasma metabolites, and metabolic syndrome traits in the METSIM cohort. Genome Biol 2017;18:1–14. doi:10.1186/s13059-017-1194-2

51 Wilmanski T, Rappaport N, Earls JC, et al. Blood metabolome predicts gut microbiome α-diversity in humans. Nat Biotechnol 2019;37. doi:10.1038/s41587-019-0233-9

52 Le Poul E, Loison C, Struyf S, et al. Functional characterization of human receptors for short chain fatty acids and their role in polymorphonuclear cell activation. J Biol Chem 2003;278:25481–9. doi:10.1074/jbc.M301403200

53 Barber MN, Risis S, Yang C, et al. Plasma lysophosphatidylcholine levels are reduced in obesity and type 2 diabetes. PLoS One 2012;7:1–12. doi:10.1371/journal.pone.0041456

54 Zuo T, Wong SH, Lam K, et al. Bacteriophage transfer during faecal microbiota transplantation in Clostridium difficile infection is associated with treatment outcome. Gut 2018;67:634–43. doi:10.1136/gutjnl-2017-313952

