## Supplementary figures and images for "Faecal virome transplantation decrease symptoms of type-2-diabetes and obesity in a murine model"

### FigureS7

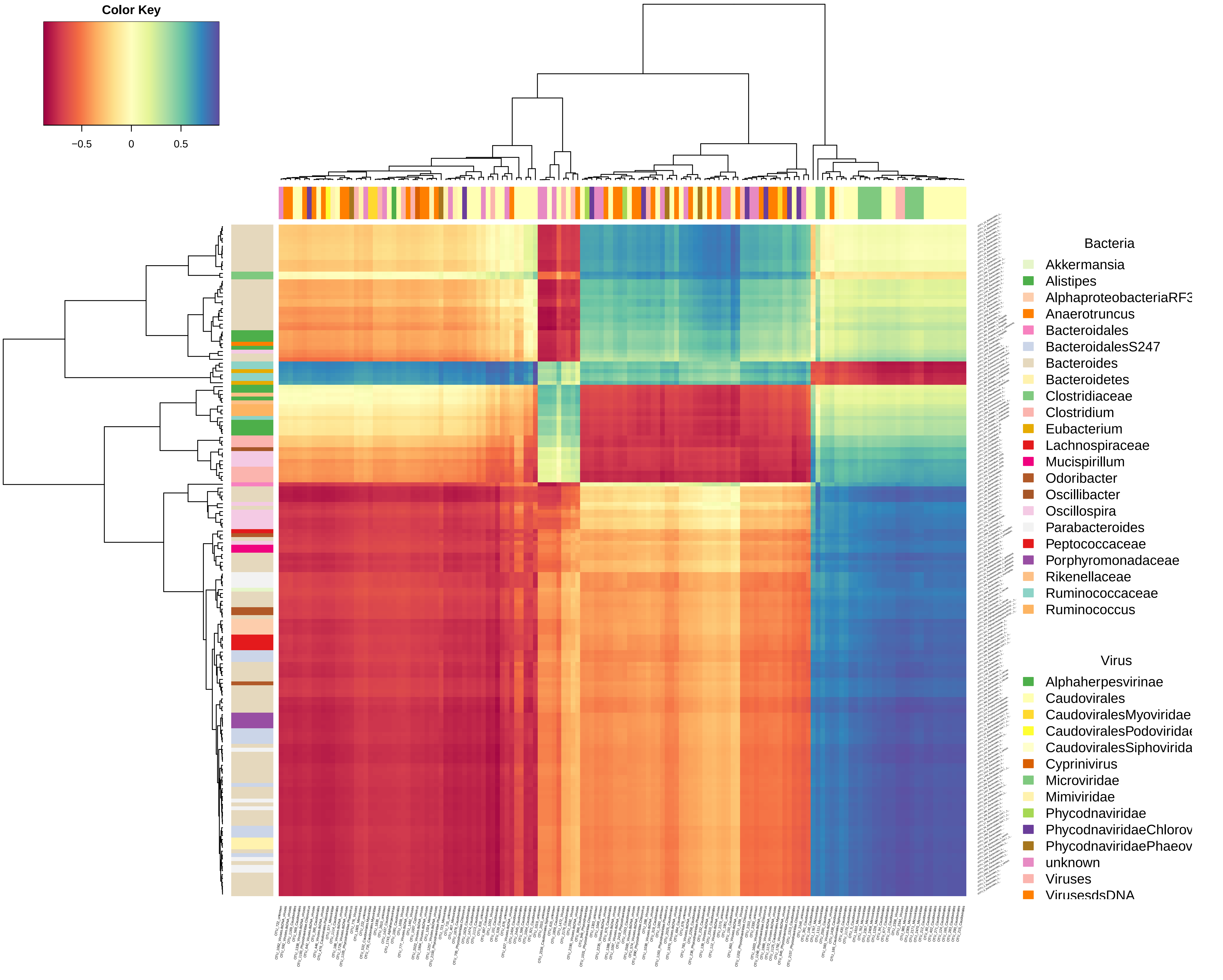

### FigureS8

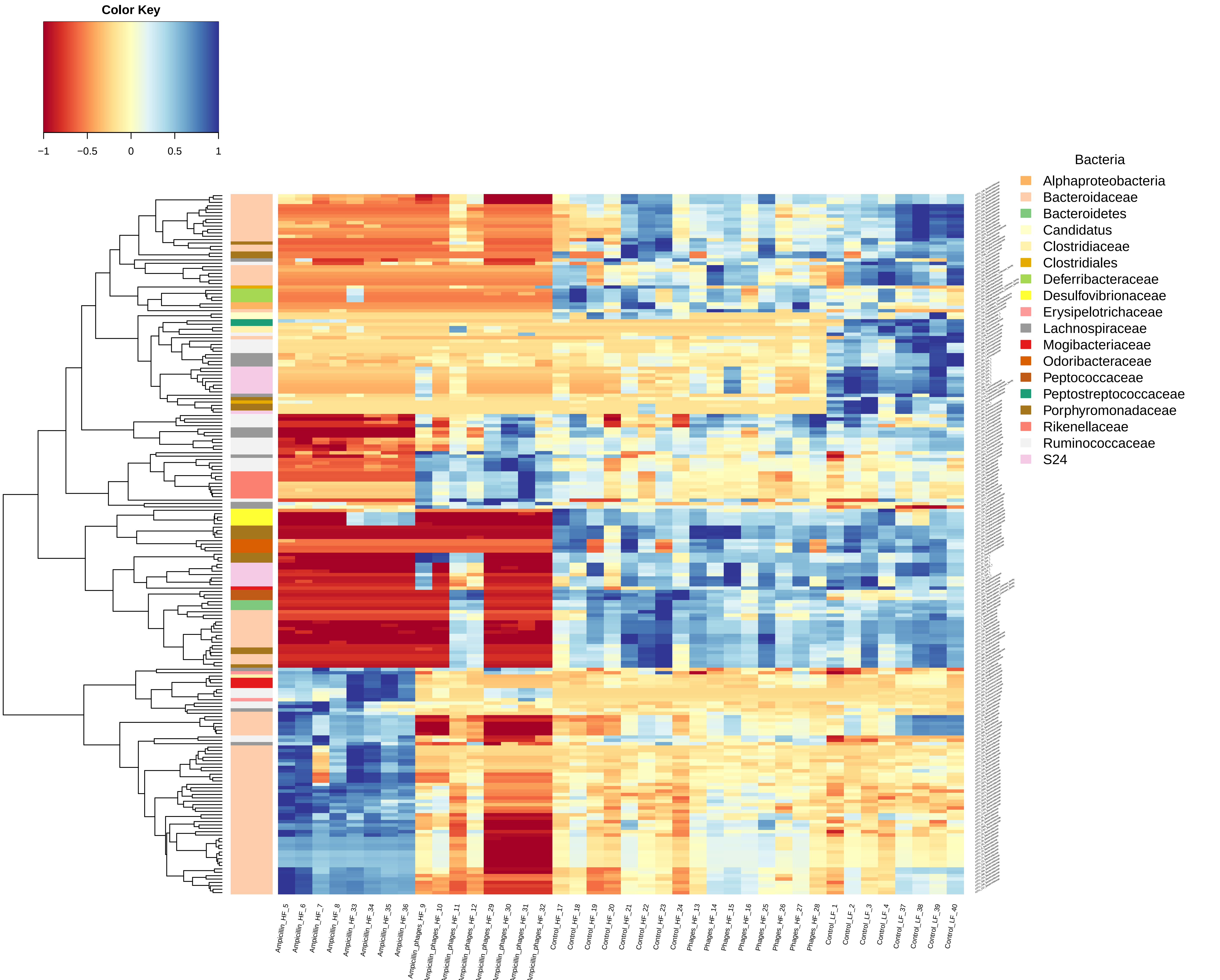

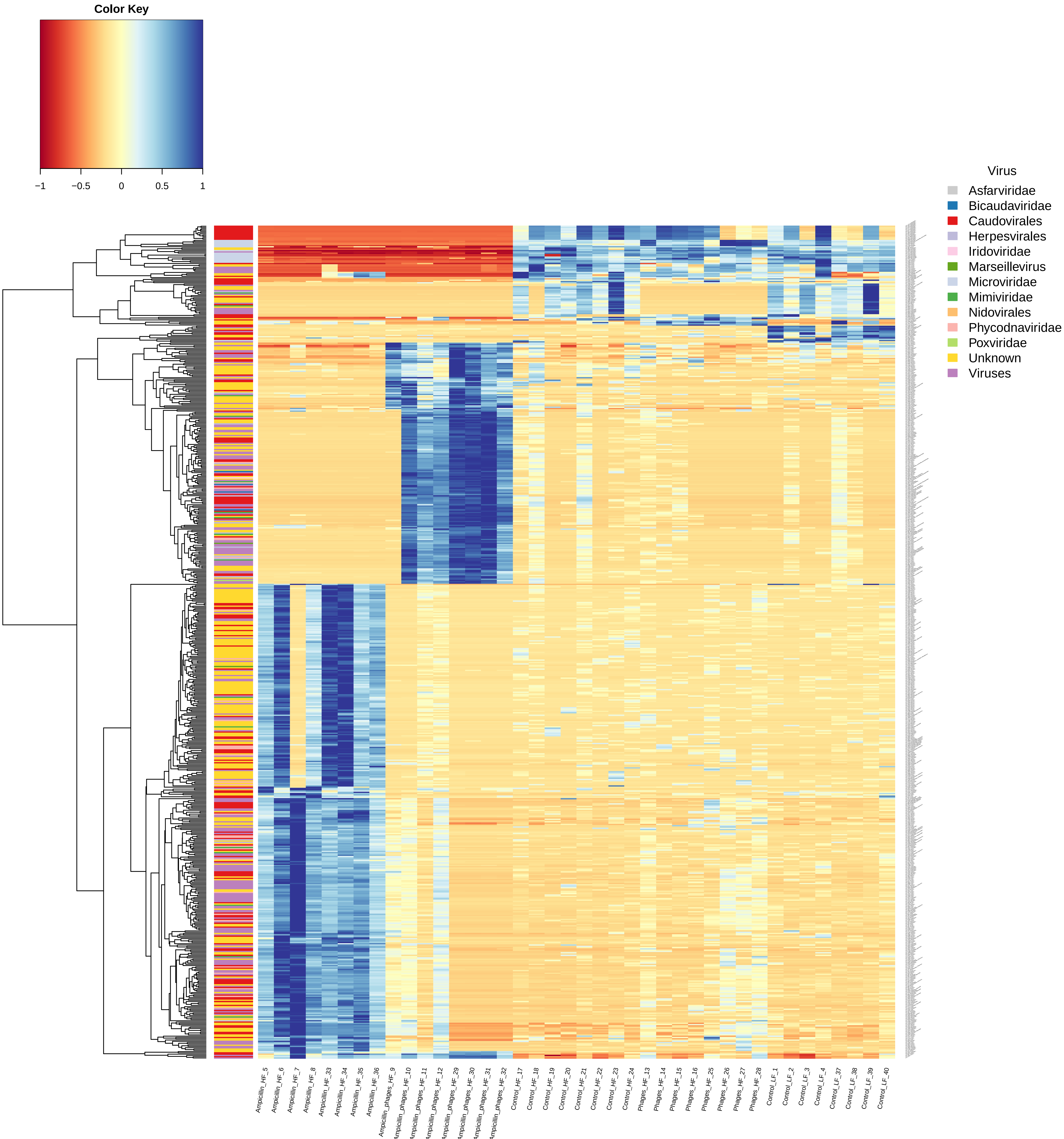
