## Supplemental materials for "Faecal virome transplantation decrease symptoms of type-2-diabetes and obesity in a murine model"

### Supplementary methods

#### Animal study design

Forty male C57BL/6NTac mice were five weeks old at arrival (Taconic Biosciences A/S, Lille Skensved, Denmark), ear tagged, enrolled into five different study groups (n=8), and housed with four animals per cage. The five groups represented: low-fat diet (LF, as lean control), high-fat diet (HF), HF+Ampicillin (Amp), HF+Amp+FVT (faecal virome transplantation) and HF+FVT. Ampicillin was included in the study, since the standard procedure of faecal microbiota transplantation (FMT) involves initial treatment with antibiotics[1]. For thirteen weeks the mice were fed *ad libitum* HF diet (Research Diets D12492, USA) or LF diet (Research Diets D12450J, USA). After six weeks on their respective diets, the HF+FVT and HF+Amp+FVT mice were treated twice with 0.15 mL FVT by oral gavage with a one-week interval (week 6 and 7) between the FVT. The FVT viromes were extracted from cecum content from eighteen mice fed a LF diet for 14 weeks, that earlier were isolated, sequenced, and analysed[2]. The titer of the FVT viromes was approx. $2 \times 10^{10}$ Virus-Like-Particles/mL (FigureS1). The remaining LF, HF+Amp, and HF mice received no treatment. For the HF+Amp and HF+Amp+FVT, the ampicillin (1 g/L ampicillin, Sigma-Aldrich A9518-25G, Schnelldorf, Germany) was added once to the drinking water one day before the FVT treatment (5 week + 6 days). Every second week, the animals were weighed, individual faeces samples were collected, non-fasted blood glucose levels were measured, and HbA1c level was measured twice (FigureS4). An oral glucose tolerance test (OGTT) was performed at 12 weeks post study (17 weeks old) on animals subjected to six hours fast[3]. Animals were orally dosed with 0.15 mL glucosemonohydrate 500 mg/ml (SAD, 823122, Sygehus Apotekerne Danmark, Denmark). Mouse data (weight, OGTT levels, etc.) were analysed in GraphPad Prism using one-way ANOVA with Tukeys test. The study was approved by the Danish Competent Authority, The Animal Experimentation Inspectorate, under the Ministry of Environment and Food of Denmark, and performed under license No. 2017-15-0201-01262 C1-3. Procedures were carried out in accordance with the Directive 2010/63/EU and to the Danish law LBK Nr 726 af 09/091993, and housing conditions as earlier described[2].

#### Termination and sampling

The mice were terminated at 18 weeks of age (13 weeks post study start) after anaesthesia with Hypnorm/midazolam mixture. Both Hypnorm (Hypnorm BN: P736/005, VetaPharma Ltd, Leeds, UK) and Midazolam were mixed with sterile water in the ratio of 1:1 (Midazolam, BN: 353 0418, Braun, Melsungen, Germany). Blood was sampled retro-orbitally and animals were euthanized by cervical dislocation. The distal half of the left lobe of liver (Lobus sinister lateralis hepatis) was sampled. Approximately 1.5 cm of the distal ileum was sampled. Ileum was dissected free from fat and connective tissue. All tissues were snap frozen in liquid nitrogen and stored in -80°C. Faecal content of caecum and colon was gently squeezed out by propulsive movement with blunt tweezers prior to snap freezing. Faecal and intestinal content was stored at room temperature during sampling and stored at -80°C. Surgical equipment used for tissue- and faecal sampling during terminal procedures was sterilised between each animal.

#### Pre-processing of faecal samples

This present study included in total 79 faecal content samples, of which 40 were isolated from cecum and 39 from colon (one mouse had no faecal content in the distant colon). Pre-processing was performed as earlier described[2]. In brief, faecal content was suspended in 1x SM buffer, homogenised, and centrifuged. The faecal supernatant was sampled for viral DNA extraction and the faecal pellet was re-suspended in 1x SM-buffer for bacterial DNA extraction.

#### Bacterial DNA extraction, sequencing, and pre-processing of raw data

Tag-encoded 16S rRNA gene amplicon sequencing was performed on an Illumina NextSeq using v2 MID output 2x150 cycles chemistry (Illumina, CA, USA). DNA extraction and library building for amplicon sequencing was performed in accordance with Krych et al. [4]. The average sequencing depth (Accession: PRJEB32560, available at ENA) for the cecum 16S rRNA gene amplicons was 164,147 reads (min. 22,732 reads and max. 200,2003 reads) and 166,012 reads for colon (min. 89,528 reads and max. 207,924 reads), see Table S1 for further details. The raw NextSeq generated dataset containing pair-ended reads were treated as earlier described[2]. In brief, the UPARSE[5] and UNOISE[6] pipeline were applied to generate high quality zOTU-tables and the SINTAX[7] algorithm was used to predict taxonomy using Ribosomal Database Project (Release 11, update 5)[8] as well as Greengenes (v13.8)[9] 16S rRNA gene collection as a reference database. The zOTU's will subsequently be referred to as bacterial OTU's (bOTU's) to differentiate from the viral counterpart. Bacterial density in the cecum and colon content was estimated by quantitative real-time polymerase chain reaction (qPCR) as previously described[10], using 16S rRNA gene primers (V3 region) as applied for the amplicon sequencing[4].

#### Viral DNA extraction, sequencing and pre-processing of raw data

The faecal supernatant was sterilely filtered through a Minisart® High Flow PES syringe filter (Cat. No. 16533, Sartorius, Germany), and was concentrated using Centriprep® Ultracel® YM-50K units (Cat. No. 4310, Millipore, USA). This constituted the concentrated virome. Viral DNA was extracted, and Illumina NextSeq sequencing data were pre-processed as previously described[2]. In brief, nuclease treatment was performed prior DNA extraction with QIAamp® Viral RNA Mini kit (Cat. No. 52904, Qiagen, Germany) that is included in the NetoVIR protocol[11], followed by 30 minutes of multiple displacement amplification (MDA) with the Illustra Ready-To-Go GenomiPhi V3 DNA Amplification Kit (Cat. No. 25-6601-96, GE Healthcare Life Sciences, UK). The MDA products was cleaned with Genomic DNA Clean & Concentrator™-10 units (Cat. No D4011, Zymo Research, USA). Library preparation was obtained with Nextera XT DNA Library Preparation Kit (Cat. No. FC-131-1096, Illumina, USA), followed by Illumina NextSeq sequencing using v2 MID output 2x150 cycles chemistry (Illumina, CA, USA). The average sequencing depth (Accession: PRJEB32560, available at ENA) for the cecum viral metagenome was 612,640 reads (min. 277,582 reads and max. 1,219,178 reads) and 356,976 reads for colon (min. 33,773 reads and max. 584,681 reads), see Table S1 for further details. The raw reads were treated with Trimmomatic[12] and Usearch[5]. Contig assembly was performed with Spades v.3.5.0[13,14], and contaminations of non-viral contigs like bacteria, human, mice, and plant DNA were removed (Figure S??). Viral contigs was identified with Kraken2[15], VirFinder[16], PHASTER[17], and virus orthologous proteins (www.vogdb.org). All contigs matching viral proteins, viral k-mers, including those that did not match any database, were subsequently retained and categorised as viral contigs. Following assembly and quality control, high-quality/dereplicated reads from all samples were merged and recruited against all the assembled contigs at 95% similarity using Subread[18]. A contingency-table of reads per Kb of contig sequence per million reads sample (RPKM) was generated with a 10x coverage threshold, here defined as the vOTU-table (viral-operational taxonomic unit). Only vOTUs ≥ 1000 bp was included in the vOTU-table.

#### Gene expression assay

Frozen liver (~30mg) and ileum (1.5 cm) pieces were transferred to tubes (FastPrep® 50-76-200, Mpbio) containing 0.6g glass beads (G4649, Sigma-Aldrich), 600 µl lysis binding solution concentrate (AM1830, Invitrogen™) and 0.7% beta-mercaptoethanol (M6250, Sigma-Aldrich).

The samples were homogenised using the FastPrep-24™ Classic Instrument (Mpbio) with 4 x (45 sec at speed 6.5) runs. The homogenate was centrifuged at 16000 x g and the supernatant was frozen at -20°C overnight. 100 µl homogenate was purified to RNA using the MagMax™ Express Magnetic Particle Processor (Applied Biosystems ™) using manufacturer's instructions (kit and protocol: AM1830, Invitrogen™). RNA purity and concentration were assessed using NanoDrop ND-1000 (NanoDrop Technologies) and RNA integrity was determined by RNA quality indicator (RQI) numbers obtained on the Experion (Biorad) platform and by 1.4% agarose gel. RQI number < 7 was accepted if the RNA had clear 18S RNA and 28S RNA bands on the gel without smearing. Mean RQI and standard deviation was 8.3 ± 0.3 for liver RNA and 8.8 ± 0.5 for ileum RNA. cDNA synthesis was performed in duplicates from 200 ng RNA as previously described[19]. Negative controls were made by excluding the reverse transcriptase in the cDNA synthesis reaction from one sample/tissue (-RT control). cDNA was diluted 1:8 and stored on -80°C before qPCR.

Genes were selected based on relevant pathways for each tissue. For measuring liver gene expression genes involved in metabolic pathways (triglyceride, carbohydrate, bile and cholesterol metabolism) as well as inflammation were selected. For ileum expression genes involved in inflammation, gut microbiota signalling and gut barrier function were selected[20,21]. Primers (Sigma-Aldrich) were designed to span an intron if possible and yield products around 75-200 nucleotides long using primer 3 (http://bioinfo.ut.ee/primer3/) or primer blast (https://www.ncbi.nlm.nih.gov/tools/primer-blast/) with standard settings[22,23]. Primer sequences are listed in TableS2. qPCR was performed using the Biomark HD system (Fluidigm Corporation) on 2x 96.96 IFC chips on pre-amplified cDNA duplicates using manufacturer's instructions with minor adjustments as previously described[19]. Melting curves were assessed prior to further analysis and primer assays yielding multiple products and/or assays yielding a product in the –RT samples were abolished. A calibration curve made from a 5x dilution row of a pool of the pre-amplified cDNA was used to calculate primer efficiency. Primer assays with efficiencies between 80-110 % were accepted for further analysis. All qPCR data processing prior to statistics was performed in Genex6 (multiD Analysis AB). 75 candidate genes and 8 reference genes were assessed in ileum and 71 candidate genes and 8 reference genes were assessed in the liver. The reference genes were analysed for stable expression using the geNorm and NormFinder algorithms[24,25]. Rplp2, Hprt, Actb, Ywhaz and Sdha were the most stable reference genes for Ileum and Actb, B2m, H2afz, Pgk1 and Sdha were most stable in liver. Candidate gene expression in cycle of quantification avalues (Cq) were normalized to the reference genes, cDNA duplicates were averaged and relative expression of the lowest expressed sample per tissue were set to 1 and the data were log2 transformed before statistical analysis in R using linear models with either HF or LF as control groups.

#### Blood plasma metabolome analysis

Plasma samples were prepared for ultra-performance liquid chromatography mass spectrometry (UPLC-MS) analysis according to a previously published protocol[26]. Briefly, prechilled methanol was added to the samples in a volume ratio 1:3. Samples were vortexed for 1 minute and left at -20°C overnight. Samples were centrifuged at 14,000 x *g* for 20 minutes at 4°C, and the supernatant was collected for analysis. Samples were analysed using an untargeted reversed phase chromatographic method previously described in the literature; a 30 minutes analytical run was used to separate and detect polar metabolites and lipids[27]. A 2.1 × 100 mm (1.7 µm) HSS T3 Acquity column (Waters Corp., Milford, MA) was used together with a Synapt G2-S QTOF mass spectrometer (Waters Corp., Manchester, UK) in both positive and negative ionization modes (ESI+ and ESI-). An imidazole propionic acid (IPPA) standard was run in both modes to assess the presence of the compound in the samples. The obtained spectra were converted into .mzML files with MSConvert and analysed with XCMS 3.6 package in R 3.6[28–31]. Briefly, peak picking was performed using the centwave algorithm and peaks were grouped with the density method. No retention time alignment was performed and features with missing values were excluded from the analysis. The remaining features were normalized using median fold change and imported into SIMCA-P 15.0 (Umetrics). Data were log transformed and pareto scaled to perform both unsupervised and supervised multivariate statistical analyses (PCA and OPLS-DA). Among the features driving the different models, only those with variable importance in the projection (VIP) scores > 2 were further investigated. Putative annotation was achieved through searching for the m/z values in online databases such as HMDB (http://www.hmdb.ca), METLIN (http://metlin.scripps.edu) and Lipidmaps (http://www.lipidmaps.org). Additionally, fragmentation patterns derived from $MS^e$ experiment were compared to online spectra when available. BLASTX v. 2.7.1[32] database was applied to annotate and evaluate the presence of known integrase genes with a minimum E-value = $10^{-3}$ and alignment length at 51 bp amongst vOTU's with a contig size above 3000 bp.

#### Bioinformatic analysis of bacterial and viral DNA

Prior any analysis the raw read counts in the vOTU-tables were normalised by reads per kilo base per million mapped reads (RPKM)[33]. B- and vOTU's which persisted in less than 8% of the samples were discarded to reduce noise, however still maintaining an average total abundance close to 98%. Cumulative sum scaling[34] (CSS) was applied for analysis of β-diversity to counteract that a few b- and vOTU's represented larger count values, and since CSS have been benchmarked with a high accuracy for the applied Bray-Curtis dissimilarity-metric[35]. CSS normalisation was executed using the Quantitative Insight Into Microbial Ecology 1.9.1[36] (QIIME 1.9.1) normalize_table.py. The viral and bacterial α-diversity analysis was based on, respectively, RPKM normalised and raw read counts to avoid bias with rarefaction[37]. QIIME 2 (2019.4)[36] plugins were used for subsequent analysis steps of α- and β-diversity statistics. The Shannon diversity index represented likewise the determined α-diversity measure. ANOSIM and Kruskal Wallis was used to evaluate multiple group comparisons. Regularized Canonical Correlation Analysis (rCCA) was performed with mixOmics v. 6.8.0. R package[38] to predict correlations between bacterial and viral taxa. Only vOTUs ≥ 5000 bp were included. The machine learning algorithm random forest[39] was applied to select variables explaining the dataset and normalised in range of -1:1 ((x-mean)/max(abs(x-mean)) and visualised by Heatmap3[40].

### Supplementary references

1 Liubakka A, Vaughn BP. Clostridium difficile Infection and Fecal Microbiota Transplant. *AACN Adv Crit Care* 2016;**27**:324–37. doi:10.4037/aacnacc2016703

2 Rasmussen TS, de Vries L, Kot W, *et al.* Mouse Vendor Influence on the Bacterial and Viral Gut Composition Exceeds the Effect of Diet. *Viruses* 2019;**11**:435. doi:10.3390/v11050435

3 Rune I, Rolin B, Lykkesfeldt J, *et al.* Long-term Western diet fed apolipoprotein E-deficient rats exhibit only modest early atherosclerotic characteristics. *Sci Rep* 2018;**8**:1–12. doi:10.1038/s41598-018-23835-z

4 Krych Ł, Kot W, Bendtsen KMB, *et al.* Have you tried spermine ? A rapid and cost-e ff ective method to eliminate dextran sodium sulfate inhibition of PCR and RT-PCR. *J Microbiol Methods J* 2018;**144**:1–7. doi:10.1016/j.mimet.2017.10.015

5 Edgar RC. UPARSE: highly accurate OTU sequences from microbial amplicon reads. *Nat Methods* 2013;**10**:996–8. doi:10.1038/nmeth.2604

6 Edgar RC. UNOISE2: improved error-correction for Illumina 16S and ITS amplicon sequencing. *bioRxiv* 2016;:081257. doi:10.1101/081257

7 Edgar R. SINTAX: a simple non-Bayesian taxonomy classifier for 16S and ITS sequences. *bioRxiv* 2016;:074161. doi:10.1101/074161

8 Cole JR, Wang Q, Fish JA, *et al.* Ribosomal Database Project: Data and tools for high throughput rRNA analysis. *Nucleic Acids Res* 2014;**42**:D633-42. doi:10.1093/nar/gkt1244

9 Mcdonald D, Price MN, Goodrich J, *et al.* An improved Greengenes taxonomy with explicit ranks for ecological and evolutionary analyses of bacteria and archaea. *ISME J* 2011;**6**:610–8. doi:10.1038/ismej.2011.139

10 Ellekilde M, Krych L, Hansen CHFHF, *et al.* Characterization of the gut microbiota in leptin deficient obese mice - Correlation to inflammatory and diabetic parameters. *Res Vet Sci* 2014;**96**:241–50. doi:10.1016/j.rvsc.2014.01.007

11 Conceição-Neto N, Zeller M, Lefrère H, *et al.* Modular approach to customise sample preparation procedures for viral metagenomics: a reproducible protocol for virome analysis. *Sci Rep* 2015;**5**. doi:10.1038/srep16532

12 Bolger AM, Lohse M, Usadel B. Trimmomatic: A flexible trimmer for Illumina sequence data. *Bioinformatics* 2014;**30**:2114–20. doi:10.1093/bioinformatics/btu170

13 Nurk S, Meleshko D, Korobeynikov A PP. metaSPAdes: A New Versatile Metagenomic Assembler. *Genome Res* 2017;**1**:30–47. doi:10.1101/gr.213959.116.4

14 Bankevich A, Nurk S, Antipov D, *et al.* SPAdes: A new genome assembly algorithm and its applications to single-cell sequencing. *J Comput Biol* 2012;**19**:455–77. doi:10.1089/cmb.2012.0021

15 Wood DE, Salzberg SL. Kraken: ultrafast metagenomic sequence classification using exact alignments. *Genome Biol* 2014;**15**:R46. doi:10.1186/gb-2014-15-3-r46

16 Ren J, Ahlgren NA, Lu YY, *et al.* VirFinder: a novel k-mer based tool for identifying viral sequences from assembled metagenomic data. *Microbiome* 2017;**5**:69. doi:10.1186/s40168-017-0283-5

17 Arndt D, Grant JR, Marcu A, *et al.* PHASTER: a better, faster version of the PHAST phage search tool. *Nucleic Acids Res* 2016;**44**:1–6. doi:10.1093/nar/gkw387

18 Liao Y, Smyth GK, Shi W. The Subread aligner: Fast, accurate and scalable read mapping by seed-and-vote. *Nucleic Acids Res* 2013;**41**. doi:10.1093/nar/gkt214

19 Mentzel CMJ, Cardoso TF, Pipper CB, *et al.* Deregulation of obesity-relevant genes is associated with progression in BMI and the amount of adipose tissue in pigs. *Mol Genet Genomics* 2018;**293**:129–36. doi:10.1007/s00438-017-1369-2

20 Bendtsen KM, Hansen CHF, Krych Ł, *et al.* Immunological effects of reduced mucosal integrity in the early life of BALB/c mice. *PLoS One* 2017;**12**:1–20. doi:10.1371/journal.pone.0176662

21 Zachariassen LF, Krych L, Rasmussen SH, *et al.* Cesarean Section Induces Microbiota-Regulated Immune Disturbances in C57BL/6 Mice. *J Immunol* 2019;**202**:142–50. doi:10.4049/jimmunol.1800666

22 Koressaar T, Remm M. Enhancements and modifications of primer design program Primer3. *Bioinformatics* 2007;**23**:1289–91. doi:10.1093/bioinformatics/btm091

23 Ye J, Coulouris G, Zaretskaya I, *et al.* Primer-BLAST: a tool to design target-specific primers for polymerase chain reaction. *BMC Bioinformatics* 2012;**13**:134.

doi:10.1186/1471-2105-13-134

24 Vandesomlele J. Accurate normalization of real-time quantitative RT-PCR data.pdf. *Genome Biol* 2002;**3**:1–12. doi:10.1186/gb-2002-3-7-research0034

25 Andersen CL, Jensen JL, Ørntoft TF. Normalization of Real-Time Quantitative Reverse Transcription-PCR Data: A Model-Based Variance Estimation Approach to Identify Genes Suited for Normalization, Applied to Bladder and Colon Cancer Data Sets. *Cancer Res* 2004;**64**:5245–50. doi:10.1158/0008-5472.CAN-04-0496

26 Sarafian MH, Gaudin M, Lewis MR, *et al.* Objective Set of Criteria for Optimization of Sample Preparation Procedures for Ultra-High Throughput Untargeted Blood Plasma Lipid Profiling by Ultra Performance Liquid Chromatography−Mass Spectrometry. *Anal Chem* 2014;**86**:5766–74. doi:10.1021/ac500317c

27 Want EJ, Coen M, Masson P, *et al.* Ultra Performance Liquid Chromatography-Mass Spectrometry Profiling of Bile Acid Metabolites in Biofluids: Application to Experimental Toxicology Studies. *J D Hepatic Transp Bile Secret Physiol Pathophysiol* 1977;**6**:5282–9. doi:10.1021/ac1007078

28 Chambers MC, Maclean B, Burke R, *et al.* A cross-platform toolkit for mass spectrometry and proteomics. *Nat Biotechnol* 2012;**30**:918–20. doi:10.1038/nbt.2377

29 Smith CA, Want EJ, O'Maille G, *et al.* XCMS:  Processing Mass Spectrometry Data for Metabolite Profiling Using Nonlinear Peak Alignment, Matching, and Identification. *Anal Chem* 2006;**78**:779–87. doi:10.1021/ac051437y

30 Tautenhahn R, Böttcher C, Neumann S. Highly sensitive feature detection for high resolution LC/MS. *BMC Bioinformatics* 2008;**9**:504. doi:10.1186/1471-2105-9-504

31 Benton HP, Want EJ, Ebbels TMD. Correction of mass calibration gaps in liquid chromatography–mass spectrometry metabolomics data. *Bioinformatics* 2010;**26**:2488–9. doi:10.1093/bioinformatics/btq441

32 Pundir S, Martin MJ, O'Donovan C. UniProt Tools. In: *Current Protocols in Bioinformatics*. Hoboken, NJ, USA: : John Wiley & Sons, Inc. 2016. 1.29.1-1.29.15. doi:10.1002/0471250953.bi0129s53

33 Roux S, Emerson JB, Eloe-Fadrosh EA, *et al.* Benchmarking viromics: an *in silico* evaluation of metagenome-enabled estimates of viral community composition and

diversity. *PeerJ* 2017;**5**:e3817. doi:10.7717/peerj.3817

34 Paulson JN, Stine OC, Bravo HC, *et al.* Differential abundance analysis for microbial marker-gene surveys. *Nat Methods* 2013;**10**:1200–2. doi:10.1038/nmeth.2658

35 Weiss S, Xu ZZ, Peddada S, *et al.* Normalization and microbial differential abundance strategies depend upon data characteristics. *Microbiome* 2017;**5**:27. doi:10.1186/s40168-017-0237-y

36 Caporaso JG, Kuczynski J, Stombaugh J, *et al.* QIIME allows analysis of high-throughput community sequencing data. *Nat Methods* 2010;**7**:335–6. doi:10.1038/nmeth.f.303

37 McMurdie PJ, Holmes S. Waste Not, Want Not: Why Rarefying Microbiome Data Is Inadmissible. *PLoS Comput Biol* 2014;**10**. doi:10.1371/journal.pcbi.1003531

38 Rohart F, Gautier B, Singh A, *et al.* mixOmics: An R package for ‘omics feature selection and multiple data integration. *PLOS Comput Biol* 2017;**13**:e1005752. doi:10.1371/journal.pcbi.1005752

39 Breiman L. Random forests. *Mach Learn* 2001;**45**:5–32. doi:10.1023/A:1010933404324

40 Zhao S, Guo Y, Sheng Q, *et al.* Heatmap3: An improved heatmap package with more powerful and convenient features. *BMC Bioinformatics* 2014;**15**:P16. doi:10.1186/1471-2105-15-S10-P16

### Supplementary tables

**Table S1.** Illumina NextSeq sequencing details of both the 16S rRNA gene amplicons and metaviromes of faecal content isolated from both cecum and colon. The table list the amount of b- and vOTUs after filtering based on 10x coverage and abundance as described in methods. Only reads used for vOTU's are included whereas reads associated for contaminations are excluded. BC = bacterial community, VC = viral community.

| | **BC** | | **VC** | |
|---|---|---|---|---|
| | **Cecum** | **Colon** | **Cecum** | **Colon** |
| Mean sequencing depth (reads) | 164147 | 166012 | 612640 | 356976 |
| STD (reads) | 31452 | 20294 | 239471 | 33773 |
| Lowest sequencing depth (reads) | 22732 | 89528 | 277582 | 33773 |
| Maximum sequencing depth (reads) | 200203 | 207924 | 1219178 | 584681 |
| vOTUs before 10x coverage filtering | - | - | 15513 | 15513 |
| bOTU/vOTUs before abundance filtering | 2968 | 2968 | 12628 | 12628 |
| bOTU/vOTUs after abundance filtering | 1515 | 1145 | 8939 | 7207 |

**TableS2:** List of primer sequences used for measuring gene expression levels in liver and ileum tissue with qPCR. All genes are included either as reference or due to known/suggested activity in metabolic syndrome. Gene names as well as related pathways are listed.

#### qPCR in liver tissue

| Gene name | Forward primer | Reverse primer | Amplicon length | Category/pathway |
|---|---|---|---|---|
| Abca1 | GCTTGTTGGCCTCAGTTAAGG | GTAGCTCAGGCGTACAGAGAT | 135 | Cholesterol Metabolism |
| Abcb11 | ACACCATTGTATGGATCAACAGC | CACCAACTCCTGCGTAGATGC | 120 | Bile salt transport |
| Abcg8 | CTGTGGAATGGGACTGTACTTC | GTTGGACTGACCACTGTAGGT | 108 | Cholesterol Metabolism |
| Actb | CGCAGCCACTGTCGAGT | CCCACGATGGAGGGGAATAC | 194 | Reference gene |
| Adipor1 | TCTTCGGGATGTTCTTCCTGG | TTTGGAAAAAGTCCGAGAGACC | 104 | Lipid metabolism |
| Adipor2 | GGCCCATCATGCTATGGAAC | GTGAGGGATCACTCGCCATC | 76 | Lipid metabolism |
| Akt1 | CATGCAGCACCGGTTCTTTG | TAGGAGAACTTGATCAGGCGG | 166 | Mtorc pathway |
| Apoa4 | AGCAGCTCAGTACCCTCTTCCA | GTACGACAAAGGGCACCAGC | 90 | Cholesterol Metabolism |
| B2m | CTGGTGCTTGTCTCACTGAC | GGTGGGTGGCGTGAGTATA | 77 | Reference gene |
| C3 | CAACACCAGCTACATCATTGG | CTGTGAATGCCCCAAGTTCT | 113 | Inflammation |
| Cd36 | AGATGACGTGGCAAAGAACAG | CCTTGGCTAGATAACGAACTCTG | 83 | Cholesterol Metabolism |
| Cd68 | TGTCTGATCTTGCTAGGACCG | GAGAGTAACGGCCTTTTTGTGA | 75 | Macrophage marker |
| Col1a1 | GGCTCCTGCTCCTCTTAGGG | TGGGGACCCTTAGGCCATTG | 96 | ECM,Liverfibrosis |
| Cpt1a | CTCCGCCTGAGCCATGAAG | CACCAGTGATGATGCCATTCT | 100 | Fatty acid oxidation |
| Csad | GGAGAGGCAGATCATTCTGGC | GCTGGCAAACATCGGCAAT | 122 | Cholesterol Metabolism |
| Cyp27a1 | ACAGGAGGGCAAGTACCCAA | AAAGCCTGACGCAGATGGTA | 126 | Bile acid synthesis |
| Cyp7a1 | GGGATTGCTGTGGTAGTGAGC | GGTATGGAATCAACCCGTTGTC | 100 | Cholesterol Metabolism |
| Cyp8b1 | CCTCTGGACAAGGGTTTTGTG | GCACCGTGAAGACATCCCC | 112 | Cholesterol Metabolism |
| Fasn | AGGTGGTGATAGCCGGTATGT | TGGGTAATCCATAGAGCCCAG | 138 | Lipogenesis |
| Fatp5 | TTGCATTCCTGTGGAGCCAG | TTTCCGATTGGACTCGGCCT | 106 | Fatty acid elongation |
| Fgf21 | GTGTCAAAGCCTCTAGGTTTCTT | GGTACACATTGTAACCGTCCTC | 123 | Glucose metabolism |
| Fgfr4 | CCTTCCACGGGGAGAATCG | CTCCACAAGGCATGTGTATGT | 113 | Glucose metabolism |
| Fxr | GCTGAGACTGGGTACCAGGG | TCGGAAGAAACCTTTGCAGCC | 183 | Bile acid |
| G6pase | CAGTGGTCGGAGACTGGTTC | GTCCAGGACCCACCAATACG | 77 | Mtorc pathway |
| Gcgr | ATGCCACCACAACCTAAGCC | GCCCACACCTCTTGAACACT | 179 | Glucose metabolism |
| Gck | AGGAGGCCAGTGTAAAGATGT | CTCCCAGGTCTAAGGAGAGAAA | 90 | Glucose metabolism |
| Ghr | CTGCAAAGAATCAATCCAAGCC | CAGTTCAGGGGAACGACACTT | 78 | Growth hormone related |
| Gusb | AAGAATACGTGGTCGGAGAGC | TCTCTGGCGAGTGAAGATCC | 107 | Reference gene |
| Gys2 | CGCTCCTTGTCGGTGACATC | CATCGGCTGTCGTTTTGGC | 160 | Glucose metabolism |
| H2afz | CTAGGACAACCAGCCACGGAC | GCGATTTGTGGATGTGTGGGAT | 168 | Reference gene |
| Hmgcl | CCGGCATCAACTACCCAGTC | GCGCTGGAAACTCTCCTCTAT | 152 | Cholesterol Metabolism |
| Hmgcr | AGAGCGAGTGCATTAGCAAAG | GATTGCCATTCCACGAGCTAT | 84 | Cholesterol Metabolism |
| Hmgcs1 | AGGCGTCTTTGCTTGTGTCT | ACACTCCAACCCTCTTCCCT | 123 | Cholesterol Metabolism |

| | | | | |
|---|---|---|---|---|
| Hmgcs1 | TGAACTGGGTCGAATCCAGC | CCTGTAGGTCTGGCATTTCCT | 97 | Cholesterol Metabolism |
| Hprt | TCAGTCAACGGGGGACATAAA | GGGGCTGTACTGCTTAACCAG | 122 | Reference gene |
| Igf1 | CACATCATGTCGTCTTCACACC | GGAAGCAACACTCATCCACAATG | 220 | Growth hormone related |
| Igfbp2 | CAGACGCTACGCTGCTATCC | CCCTCAGAGTGGTCGTCATCA | 140 | Growth hormone related |
| Il18 | AGAAAGCCGCCTCAAACCTT | GCGGTTGTACAGTGAAGTCG | 183 | Inflammation |
| Il1b | GCAACTGTTCCTGAACTCAACT | ATCTTTTGGGGTCCGTCAACT | 89 | Inflammation |
| Insig1 | GAATGTCACGCTCTTCCCCG | TGTGGTTCTCCCAGGTGACTG | 138 | Cholesterol Metabolism |
| Insig2 | TAAATCACGCCAGTGCTAAAGT | GGTGACAACGGTTGCTAAGAAAG | 155 | Cholesterol Metabolism |
| Insr | ATGGGCTTCGGGAGAGGAT | GGATGTCCATACCAGGGCAC | 121 | Glucose metabolism |
| Irs1 | GCAGCCAGAGGATCGTCAAT | CGTGAGGTCCTGGTTGTGAA | 135 | Glucose metabolism |
| Irs2 | AAGGAAGCCACAGTCGTGAA | CGTTGGTCGGAAACATGCC | 141 | Glucose metabolism |
| Klb | CAGGTATGCATGCACCAGGA | CCTTCTGATGAGGGCGGAAG | 126 | Glucose metabolism |
| Lcn2 | TCTGTCCCCACCGACCAATG | TGGCTCTCTGGCAACAGGAA | 132 | Inflammation |
| Ldlr | TGACTCAGACGAACAAGGCTG | ATCTAGGCAATCTCGGTCTCC | 118 | Cholesterol Metabolism |
| Lepr | GTCTTCGGGGATGTGAATGTC | ACCTAAGGGTGGATCGGGTTT | 158 | Leptin signalling |
| Lepra | TCTTGTGTCCTACTGCTCGGA | AAGAGTGTCCGTTCTCTTTTGGAA | 135 | Leptin signalling |
| Nr1h4 | GCTGAGACTGGGTACCAGGG | TCGGAAGAAACCTTTGCAGCC | 183 | Bile acid receptor /Fxr |
| Lipc | CTCAGCACCCGGAAACACT | CGCACTCACTATCTTCCAGATCC | 122 | Lipolysis and FA oxidation |
| Lpl | GGGAGTTTGGCTCCAGAGTTT | TGTGTCTTCAGGGGTCCTTAG | 115 | Lipolysis and FA oxidation |
| pc2 | GCCACCTACCACCGACTCA | AGCACACACCAATCCCCATTT | 135 | Glucose metabolism |
| Mttp | AGCCAGTGGGCATAGAAAATC | GGTCACTTTACAATCCCCAGAG | 110 | Cholesterol Metabolism |
| Myc | AGCGACTCTGAAGAAGAGCAAG | GATGGAGATGAGCCCGACT | 104 | Apoptosis |
| Myd88 | CTGGCCTTGTTAGACCGTGAG | ACCTGTAAAGGCTTCTCGGAC | 116 | Inflammation |
| Pck1 | AGCATTCAACGCCAGGTTC | CGAGTCTGTCAGTTCAATACCAA | 118 | Glucose metabolism |
| Pdhb | GTGGAAGAAATACGGTGACAAGA | ACCTGGTCAATAGCTTGCATAGA | 150 | Glucose metabolism |
| Pdk4 | CCGCTTAGTGAACACTCCTTC | TGACCAGCGTGTCTACAAACT | 134 | Glucose metabolism |
| Pgk1 | GGTGTTGCCAAAATGTCGCT | GGACTTGGCTCCATTGTCCA | 183 | Reference gene |
| Ppara | AACATCGAGTGTCGAATATGTGG | CCGAATAGTTCGCCGAAAGAA | 99 | Lipolysis and FA oxidation |
| Pparg | GGAAGACCACTCGCATTCCTT | GTAATCAGCAACCATTGGGTCA | 121 | Lipogenesis |
| Ppargc1a | AAGTGGTGTAGCGACCAATCG | AATGAGGGCAATCCGTCTTCA | 161 | Lipolysis and FA oxidation |
| Ptp1b | GGAACTGGGCGGCTATTTACC | CAAAAGGGCTGACATCTCGGT | 117 | Leptin related |
| Rps6kb | GGGTACTTGGTAAAGGGGGC | GCCGTGTCCTTAGCATTCCT | 130 | Mtorc pathway |
| Saa2 | GAGTCTGGGCTGCTGAGAAA | ATGGTGTCCTCGTGTCCTCT | 79 | Inflammation |
| Sdha | ATTGCTACTGGGGGCTACGG | GTCCTGGCAAGGCAAACCAG | 108 | Reference gene |
| Socs3 | ATGGTCACCCACAGCAAGTTT | TCCAGTAGAATCCGCTCTCCT | 145 | Leptin signalling |
| Sod2 | ACAATCTCAACGCCACCGAG | GCTGAAGAGCGACCTGAGTTG | 78 | Oxidative stress |
| Sqstm1 | GAACTCGCTATAAGTGCAGTGT | AGAGAAGCTATCAGAGAGGTGG | 131 | Authophagy |
| Srebf1a | TAGTCCGAAGCCGGGTGGGCGCCGG CGCCAT | GATGTCGTTCAAAACCGCTGTGTGTC CAGTTC | 106 | Lipogenesis |
| Srebp1c | ATCGGCGCGGAAGCTGTCGGGGTAG CGTC | ACTGTCTTGGTTGTTGATGAGCTGGA GCAT | 116 | Lipogenesis |

| | | | | |
|---|---|---|---|---|
| Stat3 | CACCTTGGATTGAGAGTCAAGAC | AGGAATCGGCTATATTGCTGGT | 112 | Leptin related |
| Thrb | ACACCAGCAATTACCAGAGTG | GCAGCTCGAAGGGACATGA | 125 | Glucose metabolism |
| Tlr12 | ACTGGCCTAACCAAGCTTCC | CTCATTCATGCACAGCACGG | 108 | Inflammation |
| Tlr4 | TCAGAACTTCAGTGGCTGGA | AGAGGTGGTGTAAGCCATGC | 82 | Inflammation |
| Tuba | TGTCCTGGACAGGATTCGC | CTCCATCAGCAGGGAGGTG | 115 | Reference gene |
| Ucp2 | GTGGTGGTCGGAGATACCAGA | GGGCAACATTGGGAGAAGTCC | 102 | Oxidative stress |
| Usmg5 | TGGGGTTCGGACGAAGATTG | GCCTCCATATGTGGCCAGGA | 136 | Glucose metabolism |

#### qPCR in ileum tissue

| Gene name | Forward primer | Reverse primer | Amplicon length | Category/pathway |
|---|---|---|---|---|
| Actb | CCCTAAGGCCAACCGTGAAA | CAGCCTGGATGGCTACGTAC | 83 | Reference gene |
| Arg1 | ATGGGCAACCTGTGTCCTTT | TCTACGTCTCGCAAGCCAAT | 127 | M2 Macrophages |
| B2M | CTGGTGCTTGTCTCACTGAC | GGTGGGTGGCGTGAGTATA | 77 | Reference gene |
| Ccl2 | GTCCCTGTCATGCTTCTGGG | GAGTAGCAGCAGGTGAGTGG | 104 | Cytokine |
| Ccl3 | ACCATGACACTCTGCAACCA | CAACGATGAATTGGCGTGGA | 106 | Cytokine |
| Cd38 | ACTGGAGAGCCTACCACGAA | AGTGGGGCGTAGTCTTCTCT | 179 | M1 macrophages |
| Cd8a | GGATTGGACTTCGCCTGTGA | TGGGACATTTGCAAACACGC | 130 | CD8 T cells |
| Cdh1 | GAGACCAGTTTCCTCGTCCG | AGCAGCTCTGGGTTGGATTC | 137 | Microbial interaction |
| Cldn1 | TCGACTCCTTGCTGAATCTGA | CAGCCATCCACATCTTCTGC | 159 | Gut barrier |
| Cldn2 | CGCCTTTCTCTGGACCTAGT | CTTGCTTCTTGGATCCGAGC | 192 | Gut barrier |
| Csf2 | ATGCCTGTCACGTTGAATGA | CCGTAGACCCTGCTCGAATA | 108 | Cytokine |
| Ctla4 | ATGGCTTGTCTTGGACTCCG | ACCACTGAAGGTTGGGTCAC | 137 | Tregs |
| Ctnnb1 | GAGCACATCAGGACACCCAA | CCGAGCAAGGATGTGGAGAG | 122 | b-cadhedrin, Cell adhesion |
| Cxcl1 | TGCACCCAAACCGAAGTCAT | CTCCGTTACTTGGGGACACC | 122 | Neutrophil granolucytes |
| Cxcl10 | AAGTGCTGCCGTCATTTTCT | CCTATGGCCCTCATTCTCAC | 129 | Cytokine |
| Cxcl16 | CCCAGATACCGCAGGGTACTT | TTCCCATGACCAGTTCCACA | 181 | NK/iNKT |
| Cxcr6 | TGGAACAAAGCTACTGGGCT | TCGTAGTGCCCATCGTACAG | 81 | NK/iNKT |
| Defa3 | AAAACTGAGGAGCAGCCAGG | CAGCGACAGCAGAGTGTGTA | 194 | Neutrophil granolucytes |
| Defa5 | AGGCCCAGGCTGATCCTATC | CTGCAGGTCCCAAAAACGC | 199 | a-defensin , paneth cells |
| Ffar2 | AGGGAGGAATCACAGGAAACG | CTTGGGCAAGTTCAGGGGTT | 165 | Short chain fatty acid |
| Ffar3 | CAGAGTGCCAGTTGTCCAAT | GCAAAAGTAAGTCCACAGCCA | 188 | Free fatty acids receptor |
| Fgf15 | ACGGCAAGATATACGGGCTG | GGCTTGGCCTGGATGAAGAT | 133 | GM signalling |
| Foxp3 | CTCCAGGACAGACCACACTTC | TGATCATGGCTGGGTTGTC | 103 | Tregs;CD4 |
| Foxp3 | CACCTGGAAGAATGCCATC | GTCCACACTGCTCCCTTCTC | 84 | Tregs;CD4 |
| Fxr | GCTGAGACTGGGTACCAGGG | TCGGAAGAAACCTTTGCAGCC | 183 | GM signalling |
| Gata3 | GCTACGGTGCAGAGGTATCC | CAGAGATCCGTGCAGCAGAG | 75 | ILC;CD4; Th2 |
| GCG | TCTACACCTGTTCGCAGCTC | GTCCTCATGCGCTTCTGTCT | 172 | Obesity |
| Gusb | AGTATGGAGCAGACGCAATCC | ACAGCCTTCTGGTACTCCTCA | 76 | Reference gene |
| Gzmb | AGAGGGGGTACAAGGTCACA | CATGTCCCCCGATGATCTCC | 187 | CD8 T cells |
| Hprt | TCAGTCAACGGGGGACATAAA | GGGGCTGTACTGCTTAACCAG | 122 | Reference gene |
| Icam1 | CTGTGCTTTGAGAACTGTGGC | CAGGGTGAGGTCCTTGCCTA | 129 | Cell adhesion |
| Ifnγ | GGCACAGTCATTGAAAGCCTA | GCCAGTTCCTCCAGATATCCA | 103 | Innate; NK/iNKT ; Neutrophil |
| Il10 | AAAGGACCAGCTGGACAACA | TAAGGCTTGGCAACCCAAGTA | 79 | Tregs |
| Il12a | AAACCAGCACATTGAAGACC | GGAAGAAGTCTCTCTAGTAGCC | 80 | Cytokine |
| Il15 | ACAGCTCAGAGAGAATCCACC | ATGAGCTGGCTATGGCGATG | 187 | NK/iNKT |
| Il18 | GGCTGTGACCCTCTCTGTG | TGGATCCATTTCCTCAAAGG | 85 | Th1 |
| Il1b | GCAACTGTTCCTGAACTCAACT | ATCTTTTGGGGTCCGTCAACT | 89 | M1 Macrophages; Th1 |

| | | | | |
|---|---|---|---|---|
| Il1b | TGCAGCTGGAGAGTGTGG | TCAAACTCCACTTTGCTCTTGA | 100 | M1 Macrophages; Th1 |
| Il33 | GGGCTCACTGCAGGAAAGTA | TTTGCCGGGGAAATCTTGGA | 115 | Cytokine |
| Il4 | CCTGGATTCATCGATAAGCTG | TCCATTTGCATGATGCTCTT | 93 | M2 Macrophages; Th2 |
| Il6 | CAAAGCCAGAGTCCTTCAGAG | GAGCATTGGAAATTGGGGTA | 107 | Th1;Th2 |
| Irf3 | CCACAAGGACAAGGACGGAG | CCACATTTCCCCCATGCAGA | 124 | Microbial interaction |
| Itgax_v2 | CAAGACAGGACATCGCTCCC | GTGAACAGTTGGTGACACTCT | 200 | Cd11c;dendritic cells |
| Klf4 | CGAGAAACCTTACCACTGTGACT | CCTGTGTGTTTGCGGTAGTG | 90 | Transcription factor |
| Muc2 | TATGCCAGGCCAGGAGTTTA | GCAAGGCAGGTCTTTACACA | 82 | Gut barrier |
| Myc | AGCGACTCTGAAGAAGAGCAAG | GATGGAGATGAGCCCGACT | 104 | Transcription factor |
| Myd88 | CCAGGTGTCCAACAGAAGC | CTTGGTGCAAGGGTTGGTAT | 114 | Tlr signalling |
| Nfkb | GGCAGGTATTTGACATACTAAATGG | TGCAGAGTTGTAGCCTCGTG | 117 | Inflammation |
| Nfkbia | GAGCGAGGATGAGGAGAGCTA | GGCCTCCAAACACACAGTCA | 83 | Inflammation |
| Nkap | ATGTTCCGGTTACGTAATGAGTG | ATGCAAGGGCTCTCTTCTCA | 107 | Inflammation |
| Nos2 | GTGACCATGGAGCATCCCAA | TCGAACTCCAATCTCGGTGC | 159 | M1 Macrophages |
| Ocld | GCTGCTGCTGATGAATATAATAGACT | TCCCACCATCCTCTTGATGT | 120 | Gut barrier |
| Pla2g2a | GGGGCCAAATCACCTGTTCT | GTTCCGGGCGAAACATTCAG | 92 | Microbial interaction |
| Prf1 | ACACAGTAGAGTGTCGCATGT | GCCGTGATAAAGTGCGTGC | 163 | CD8 T cells |
| Pyy | GCTTCTCCCACCTTCCATCT | AGACAGGCGAGCAGGATTAG | 121 | Obesity |
| Reg3a | TGGGCTCCATGATCCAACAA | CTGTCAGACTCCCACAGTGAC | 140 | Gut barrier |
| Reg3γ | ACAGACAAGATGCTTCCCCG | AGCTGCTACGTGAAGATGGG | 127 | Gut barrier |
| Retnlb | CTGTCCTGCTGGGATGGT | CCAGTCCATGACTGAGCACT | 109 | Gut barrier |
| Rorc | TACCCTACTGAGGAGGACAGG | AATGGGGCAGTTCTGCTGAC | 200 | Th17; CD4 |
| Rplp2 | CTCCTCTCCTAGCGCCAAAG | CAACACCCTGAGCGATGACA | 134 | Reference gene |
| Sdha | ATTGCTACTGGGGGCTACGG | GTCCTGGCAAGGCAAACCAG | 108 | Reference gene |
| Smad4 | GCTCCAGCCATCAGTCTGTC | CAGCCCTTCACAAAGCTCAT | 92 | Inflammation |
| Stat4 | GAAGTACCTCTACCCTGACATTCC | AGGGGACGTAACCCTTGTCT | 113 | Th1 |
| Stat5 | GGTCCCTGAGTTCGTCAATG | GGTTGGGTGGGTACATGTTG | 116 | Inflammation |
| Tbp | ACCTTATGCTCAGGGCTTGG | TGCCGTAAGGCATCATTGGA | 83 | Reference gene |
| Tbx21_T bet | GGGCTTCCAACAATGTGACC | AGCTGAGTGATCTCTGCGTTC | 193 | ILC; Th1; CD4 |
| Tff3 | CTGTCACATCGGAGCAGTGT | CAGGGCACATTTGGGATACT | 67 | Gut barrier |
| Tgfb1 | CAACTATTGCTTCAGCTCCACA | ACTTCCAACCCAGGTCCTTC | 83 | Cytokine |
| Timp1 | GGGGTGTGCACAGTGTTTC | GACCTGATCCGTCCACAAAC | 81 | Gut barrier |
| Tjp1 | GAGATGTTTATGCGGACGGTG | CTGTTTCCTCCATTGCTGTGC | 144 | Gut barrier |
| Tl1a_Tnf sf15 | CCATCCTCGCAGGACTTAGC | TGCCTCTGGGAGGTGAGTAA | 135 | Th1 |
| Tlr11 | CCACCACCCCATGCTCAAAG | TGCCCAGCCAGTCAAGGTAA | 115 | Tlr signalling |
| Tlr12 | ACTGGCCTAACCAAGCTTCC | CTCATTCATGCACAGCACGG | 108 | Tlr signalling |
| Tlr13 | CTGCTTCCTCTGTTGCATGA | CATGTTCAAAGGCACGGTCT | 141 | Tlr signalling |
| Tlr2 | GCATCCGAATTGCATCACCG | ACAGCGTTTGCTGAAGAGGA | 136 | Tlr signalling |
| Tlr3 | GAATCACAATCGCGCACCAA | CCATAGGGACAAAAGTCCCCC | 178 | Tlr signalling |

| | | | | |
|---|---|---|---|---|
| Tlr4 | CTCTCATGGCCTCCACTGGT | TTAGGAACTACCTCTATGCAGGGAT | 137 | Tlr signalling |
| Tlr5 | GATGGATGCTGAGTTCCCCC | AAAGGCTATCCTGCCGTCTG | 139 | Tlr signalling |
| Tlr9 | GTTTGTCAGAGGGAGCCTCG | AGGCTTCAGCTCACAGGGTA | 159 | Tlr signalling |
| Tnfa | CAAATGGCCTCCCTCTCATCA | TGGGCTACAGGCTTGTCAC | 88 | Innate; M1 Macrophages |
| Tslp | TGAAACTGAGAGAAATGACGGTACT | TCTGGAGATTGCATGAAGGA | 114 | Cytokine |
| Ywhaz | GAAAAGTTCTTGATCCCCAATGC | TGTGACTGGTCCACAATTCCTT | 134 | Reference gene |
| Zbtb16 | GCACTACAGGGTTCACACAGG | CACCGTTGTGTGTTCTCAGG | 107 | NK; ILC |

**TableS3:** List of genes in liver and ileum tissue observed with significant ($p < 0.05$) inter-group differences in gene expressions when using the LF and HF group as reference level. 57 of 74 for liver tissue and 58 of 74 for ileum tissue. The remaining genes had no significant differences. The statistics are based on one-way ANOVA. HF = high-fat, LF = low-fat, FVT = faecal virome transplantation, Amp = Ampicillin.

| **Liver** | | | **Ileum** | | |
|---|---|---|---|---|---|
| **Gene** | **Groups** | **p-value** | **Gene** | **Groups** | **p-value** |
| Abca1 | | | Ccl3 | | |
| | LF vs. HF+Amp | 0.02112440000 | | HF vs. HF+Amp | 0.00212424900 |
| | LF vs. HF+Amp+FVT | 0.01620369000 | | LF vs. HF+Amp | 0.02489124000 |
| | LF vs. HF+FVT | 0.04120160000 | Cd38 | | |
| Abcb11 | | | | HF vs. HF+Amp | 0.00026688020 |
| | HF vs. LF | 0.00005377600 | | HF vs. HF+Amp+FVT | 0.01303531000 |
| | LF vs. HF+Amp | 0.00005494583 | | LF vs. HF+Amp | 0.00037004770 |
| | LF vs. HF+Amp+FVT | 0.00003656354 | | LF vs. HF+Amp+FVT | 0.01712216000 |
| | LF vs. HF+FVT | 0.00000590261 | Cd8 | | |
| Abcg8 | | | | HF vs. HF+Amp | 0.00000023656 |
| | HF vs. LF | 0.00625214900 | | HF vs. HF+Amp+FVT | 0.00000005266 |
| | LF vs. HF+Amp | 0.00023705720 | | HF vs. LF | 0.00000000231 |
| | LF vs. HF+Amp+FVT | 0.00105338600 | | LF vs. HF+FVT | 0.00000000055 |
| | LF vs. HF+FVT | 0.00382718000 | Cdh1 | | |
| Adipor1 | | | | HF vs. HF+Amp | 0.02290421000 |
| | HF vs. LF | 0.00101718800 | Csf2 | | |
| | LF vs. HF+Amp | 0.00919870400 | | LF vs. HF+Amp | 0.03345960000 |
| | LF vs. HF+Amp+FVT | 0.01855762000 | | LF vs. HF+Amp+FVT | 0.00513387500 |
| | LF vs. HF+FVT | 0.00443295100 | Ctla4 | | |
| Apoa4 | | | | LF vs. HF+Amp | 0.00004159881 |
| | HF vs. HF+Amp | 0.04302623000 | | LF vs. HF+Amp+FVT | 0.02277963000 |
| | HF vs. LF | 0.01158187000 | Ctnnb1 | | |
| | LF vs. HF+FVT | 0.04427187700 | | HF vs. HF+Amp+FVT | 0.01372074000 |
| C3 | | | | LF vs. HF+Amp+FVT | 0.02387554000 |
| | HF vs. LF | 0.02870979000 | Cxcl10 | | |
| | LF vs. HF+FVT | 0.01675755000 | | HF vs. HF+Amp | 0.00003377237 |
| Cd36 | | | | HF vs. HF+Amp+FVT | 0.00005907369 |
| | HF vs. LF | 0.00000000089 | | HF vs. LF | 0.00895956600 |
| | LF vs. HF+Amp | 0.00000003023 | | LF vs. HF+FVT | 0.00705519400 |
| | LF vs. HF+Amp+FVT | 0.00000024737 | Cxcl16 | | |
| | LF vs. HF+FVT | 0.00000000048 | | HF vs. LF | 0.00013989960 |
| Cd68 | | | | LF vs. HF+Amp | 0.00250048200 |
| | LF vs. HF+FVT | 0.00326076800 | | LF vs. HF+FVT | 0.00010916910 |
| Col1a1 | | | | LF vs. HF+Amp+FVT | 0.00358833700 |
| | HF vs. LF | 0.00280927400 | Cxcr6 | | |
| | LF vs. HF+Amp | 0.09661015210 | | HF vs. HF+Amp | 0.00005685717 |

| | | |
|---|---|---|
| | LF vs. HF+Amp+FVT | 0.00664671160 |
| | LF vs. HF+FVT | 0.00060760030 |
| Cpt1a | | |
| | HF vs. LF | 0.00000201879 |
| | LF vs. HF+Amp | 0.00001051250 |
| | LF vs. HF+Amp+FVT | 0.00003056221 |
| | LF vs. HF+FVT | 0.00000965348 |
| Csad | | |
| | HF vs. HF+Amp | 0.01542988470 |
| | HF vs. HF+Amp+FVT | 0.01907177180 |
| | LF vs. HF+Amp | 0.01316900730 |
| | LF vs. HF+Amp+FVT | 0.01632249730 |
| Cyp27a1 | | |
| | HF vs. LF | 0.00050095720 |
| | LF vs. HF+Amp | 0.03421026000 |
| | LF vs. HF+Amp+FVT | 0.00654041300 |
| | LF vs. HF+FVT | 0.00030501840 |
| Cyp7a1 | | |
| | LF vs. HF+Amp | 0.00094624370 |
| | LF vs. HF+Amp+FVT | 0.00260621800 |
| | LF vs. HF+FVT | 0.02698069000 |
| Cyp8b1 | | |
| | HF vs. LF | 0.00012299550 |
| | LF vs. HF+Amp | 0.00000066351 |
| | LF vs. HF+Amp+FVT | 0.00000176970 |
| | LF vs. HF+FVT | 0.00009409759 |
| Fasn | | |
| | HF vs. LF | 0.04569408000 |
| | LF vs. HF+Amp | 0.00026198120 |
| | LF vs. HF+Amp+FVT | 0.00144100400 |
| Fatp5 | | |
| | HF vs. LF | 0.04142051000 |
| | LF vs. HF+Amp+FVT | 0.01798626000 |
| Fgf21 | | |
| | HF vs. LF | 0.01261544000 |
| | LF vs. HF+FVT | 0.02387727000 |
| Fgfr4 | | |
| | HF vs. LF | 0.00017870680 |
| | LF vs. HF+Amp | 0.00840338300 |
| | LF vs. HF+Amp+FVT | 0.00966313300 |
| | LF vs. HF+FVT | 0.00014780350 |
| Fxr | | |
| | HF vs. LF | 0.03236542000 |
| | HF vs. HF+Amp+FVT | 0.00000040113 |
| | HF vs. LF | 0.00000024627 |
| | LF vs. HF+FVT | 0.00000000294 |
| Defa | | |
| | HF vs. LF | 0.00892109500 |
| | LF vs. HF+Amp | 0.01341588000 |
| | LF vs. HF+FVT | 0.00053602870 |
| Defa5 | | |
| | LF vs. HF+Amp | 0.00407840200 |
| | LF vs. HF+FVT | 0.03252443000 |
| Ffar2 | | |
| | HF vs. HF+Amp | 0.00039318780 |
| | HF vs. HF+FVT | 0.01360030000 |
| | HF vs. HF+Amp+FVT | 0.00227533100 |
| | HF vs. LF | 0.03007936000 |
| Ffar3 | | |
| | HF vs. LF | 0.01832678210 |
| | LF vs. HF+Amp | 0.00679956470 |
| | LF vs. HF+FVT | 0.02209521290 |
| | LF vs. HF+Amp+FVT | 0.00111273440 |
| Fgf15 | | |
| | HF vs. HF+Amp | 0.00010368100 |
| | LF vs. HF+Amp | 0.00323260100 |
| Foxp3_v1 | | |
| | LF vs. HF+Amp | 0.04835099000 |
| Foxp3_v2 | | |
| | HF vs. HF+Amp | 0.03528808000 |
| | HF vs. HF+Amp+FVT | 0.00110818000 |
| | HF vs. LF | 0.03575527000 |
| Gata3 | | |
| | HF vs. HF+Amp | 0.01462031000 |
| | HF vs. LF | 0.04300743000 |
| | LF vs. HF+FVT | 0.00051883430 |
| GCG | | |
| | LF vs. HF+Amp+FVT | 0.00701069470 |
| Gzmb | | |
| | HF vs. HF+Amp | 0.00000009253 |
| | HF vs. HF+Amp+FVT | 0.00000018309 |
| | HF vs. LF | 0.00000000762 |
| | LF vs. HF+FVT | 0.00000033505 |
| Icam1 | | |
| | LF vs. HF+Amp | 0.02480636000 |
| | LF vs. HF+Amp+FVT | 0.01862641000 |

| | | |
|---|---|---|
| | LF vs. HF+Amp | 0.00733619300 |
| | LF vs. HF+Amp+FVT | 0.03420143000 |
| | LF vs. HF+FVT | 0.00749734300 |
| G6pase | | |
| | HF vs. HF+Amp | 0.01021773000 |
| | HF vs. HF+Amp+FVT | 0.00996132700 |
| | HF vs. LF | 0.00000185390 |
| | LF vs. HF+Amp | 0.00499680100 |
| | LF vs. HF+Amp+FVT | 0.00513129500 |
| | LF vs. HF+FVT | 0.00031808980 |
| Gck | | |
| | HF vs. LF | 0.04207954000 |
| | LF vs. HF+Amp+FVT | 0.04743183000 |
| | LF vs. HF+FVT | 0.02424363000 |
| Hmgcl | | |
| | HF vs. HF+Amp+FVT | 0.02817423000 |
| | HF vs. LF | 0.00000000013 |
| | LF vs. HF+Amp | 0.00000000034 |
| | LF vs. HF+Amp+FVT | 0.00000009601 |
| | LF vs. HF+FVT | 0.00000000001 |
| Hmgcs1_v1 | | |
| | HF vs. HF+Amp | 0.02365737000 |
| | LF vs. HF+Amp | 0.00030356090 |
| | LF vs. HF+Amp+FVT | 0.00673018430 |
| | LF vs. HF+FVT | 0.00742677210 |
| Hmgcs1_v2 | | |
| | HF vs. HF+Amp | 0.02669275000 |
| | LF vs. HF+Amp | 0.00073890290 |
| | LF vs. HF+Amp+FVT | 0.01222065430 |
| | LF vs. HF+FVT | 0.01439603330 |
| Igf1 | | |
| | HF vs. LF | 0.03969776000 |
| | LF vs. HF+FVT | 0.00550111400 |
| Igfbp2 | | |
| | HF vs. HF+FVT | 0.02565159000 |
| | HF vs. LF | 0.00000002372 |
| | LF vs. HF+Amp | 0.00000051022 |
| | LF vs. HF+Amp+FVT | 0.00000129862 |
| | LF vs. HF+FVT | 0.00002678073 |
| Il18 | | |
| | LF vs. HF+Amp+FVT | 0.04756932000 |
| Il1b | | |
| | HF vs. LF | 0.00081323980 |
| Ifng | | |
| | HF vs. HF+Amp | 0.00188563700 |
| | HF vs. HF+Amp+FVT | 0.00083950170 |
| | HF vs. LF | 0.00052510100 |
| | LF vs. HF+FVT | 0.00000436080 |
| Il10 | | |
| | HF vs. HF+Amp | 0.01771579000 |
| | LF vs. HF+Amp | 0.02591173000 |
| Il15 | | |
| | HF vs. HF+Amp | 0.00128816000 |
| | HF vs. HF+Amp+FVT | 0.02263851000 |
| | HF vs. LF | 0.00227216400 |
| | LF vs. HF+FVT | 0.01639969000 |
| Il18 | | |
| | HF vs. LF | 0.00082993370 |
| | LF vs. HF+Amp | 0.00001150377 |
| | LF vs. HF+FVT | 0.00017129680 |
| | LF vs. HF+Amp+FVT | 0.00016570730 |
| Il1b_v1 | | |
| | HF vs. HF+Amp | 0.00932286300 |
| | LF vs. HF+Amp | 0.04049308000 |
| Il1b_v2 | | |
| | HF vs. HF+Amp | 0.01204419000 |
| | LF vs. HF+Amp | 0.02328293000 |
| Il23 | | |
| | HF vs. HF+Amp+FVT | 0.01519148800 |
| | LF vs. HF+Amp+FVT | 0.00496833400 |
| Il6 | | |
| | HF vs. HF+Amp+FVT | 0.03064869000 |
| Muc2 | | |
| | HF vs. LF | 0.00773860800 |
| | LF vs. HF+Amp | 0.02839686000 |
| | LF vs. HF+FVT | 0.00118373600 |
| Nfkbia | | |
| | LF vs. HF+Amp | 0.00352165200 |
| | LF vs. HF+Amp+FVT | 0.00110789500 |
| Nkap | | |
| | HF vs. HF+Amp+FVT | 0.01988487230 |
| | LF vs. HF+Amp+FVT | 0.00524404100 |
| Nos2 | | |
| | HF vs. HF+Amp | 0.00000004191 |
| | HF vs. HF+Amp+FVT | 0.00000121583 |
| | LF vs. HF+Amp | 0.00000003637 |

| | | |
|---|---|---|
| | LF vs. HF+Amp | 0.01566679000 |
| | LF vs. HF+Amp+FVT | 0.00762913500 |
| Insig1 | | |
| | HF vs. LF | 0.01254585000 |
| | LF vs. HF+Amp | 0.03288418000 |
| | LF vs. HF+Amp+FVT | 0.01956697000 |
| | LF vs. HF+FVT | 0.02866000000 |
| Insr | | |
| | HF vs. LF | 0.03701851000 |
| Irs2 | | |
| | HF vs. LF | 0.00000165550 |
| | LF vs. HF+Amp | 0.00003402056 |
| | LF vs. HF+Amp+FVT | 0.00002352490 |
| | LF vs. HF+FVT | 0.00005977001 |
| Klb | | |
| | HF vs. HF+FVT | 0.04678326000 |
| | HF vs. LF | 0.00000107070 |
| | LF vs. HF+Amp | 0.00004971667 |
| | LF vs. HF+Amp+FVT | 0.00006725277 |
| | LF vs. HF+FVT | 0.00050737590 |
| Lepr | | |
| | HF vs. HF+FVT | 0.01000781000 |
| | HF vs. LF | 0.00000088206 |
| | LF vs. HF+Amp | 0.00000064120 |
| | LF vs. HF+Amp+FVT | 0.00000344162 |
| | LF vs. HF+FVT | 0.00267448200 |
| Lepra | | |
| | HF vs. HF+FVT | 0.03667262000 |
| | HF vs. LF | 0.00000874480 |
| | LF vs. HF+Amp | 0.00000436538 |
| | LF vs. HF+Amp+FVT | 0.00001042244 |
| | LF vs. HF+FVT | 0.00460029300 |
| Lipc | | |
| | HF vs. LF | 0.00544776800 |
| | LF vs. HF+Amp | 0.03913227000 |
| | LF vs. HF+Amp+FVT | 0.01079691000 |
| | LF vs. HF+FVT | 0.00320559200 |
| Lpl | | |
| | HF vs. LF | 0.00657596800 |
| | LF vs. HF+Amp | 0.00249820100 |
| | LF vs. HF+Amp+FVT | 0.00242556400 |
| Myc | | |
| | HF vs. HF+Amp | 0.00083185070 |
| | LF vs. HF+Amp+FVT | 0.00000105211 |
| Pla2g2a | | |
| | HF vs. HF+Amp | 0.02090997000 |
| | HF vs. LF | 0.00020106910 |
| | LF vs. HF+Amp | 0.00000013804 |
| | LF vs. HF+FVT | 0.00031164030 |
| | LF vs. HF+Amp+FVT | 0.00023809810 |
| Prf1 | | |
| | HF vs. HF+Amp | 0.00622755300 |
| | HF vs. HF+Amp+FVT | 0.00024387920 |
| | HF vs. LF | 0.00597102100 |
| | LF vs. HF+FVT | 0.01554797000 |
| Pyy | | |
| | HF vs. LF | 0.00812582720 |
| | LF vs. HF+Amp | 0.01653451310 |
| | LF vs. HF+FVT | 0.02580675910 |
| Reg3g | | |
| | HF vs. HF+Amp | 0.00333428600 |
| | HF vs. HF+Amp+FVT | 0.01274441000 |
| | LF vs. HF+Amp | 0.00389322700 |
| | LF vs. HF+Amp+FVT | 0.01469302000 |
| Reg3a | | |
| | HF vs. HF+Amp | 0.00002384444 |
| | HF vs. HF+FVT | 0.01765701000 |
| | HF vs. HF+Amp+FVT | 0.00003977341 |
| | LF vs. HF+Amp | 0.00104000500 |
| | LF vs. HF+FVT | 0.00058389200 |
| | LF vs. HF+Amp+FVT | 0.00166650500 |
| Retnlb | | |
| | HF vs. HF+Amp+FVT | 0.01071542000 |
| Smad4 | | |
| | LF vs. HF+FVT | 0.02066207000 |
| Stat4 | | |
| | HF vs. HF+Amp | 0.03881425000 |
| | HF vs. HF+Amp+FVT | 0.01429195000 |
| | LF vs. HF+Amp+FVT | 0.04657572000 |
| Stat5 | | |
| | HF vs. LF | 0.00914558400 |
| | LF vs. HF+Amp | 0.04811964000 |
| Tbx21_Tbet | | |
| | HF vs. HF+Amp | 0.00098546450 |
| | HF vs. HF+Amp+FVT | 0.00008190609 |
| | HF vs. LF | 0.01102207000 |

| | | |
|---|---|---|
| | HF vs. HF+Amp+FVT | 0.00077553920 |
| | HF vs. HF+FVT | 0.04577295000 |
| | HF vs. LF | 0.00524098100 |
| Nr1h4 | | |
| | HF vs. LF | 0.03800112000 |
| | LF vs. HF+Amp | 0.00427563500 |
| | LF vs. HF+Amp+FVT | 0.03714705000 |
| | LF vs. HF+FVT | 0.00726939600 |
| Pck1 | | |
| | HF vs. LF | 0.00789564400 |
| Pdhb | | |
| | HF vs. HF+Amp | 0.00815967700 |
| Ppara | | |
| | HF vs. LF | 0.00001006209 |
| | LF vs. HF+Amp | 0.00010803110 |
| | LF vs. HF+Amp+FVT | 0.00039806240 |
| | LF vs. HF+FVT | 0.00007740611 |
| Pparg | | |
| | HF vs. LF | 0.00000000000 |
| | LF vs. HF+Amp | 0.00000000000 |
| | LF vs. HF+Amp+FVT | 0.00000000001 |
| | LF vs. HF+FVT | 0.00000000000 |
| Ppargc1a | | |
| | HF vs. HF+FVT | 0.00378615200 |
| | HF vs. LF | 0.00000030326 |
| | HF vs. LF | 0.00000030326 |
| | LF vs. HF+Amp | 0.00000164149 |
| | LF vs. HF+Amp+FVT | 0.00000006990 |
| | LF vs. HF+FVT | 0.00286712500 |
| Rps6kb | | |
| | HF vs. LF | 0.00597899300 |
| | LF vs. HF+Amp+FVT | 0.00515070500 |
| | LF vs. HF+FVT | 0.03177196000 |
| Socs3 | | |
| | HF vs. HF+Amp | 0.00079424980 |
| | HF vs. HF+Amp+FVT | 0.00059730100 |
| | HF vs. HF+FVT | 0.00129172800 |
| | LF vs. HF+Amp | 0.00097337660 |
| | LF vs. HF+Amp+FVT | 0.00073337160 |
| | LF vs. HF+FVT | 0.00157747900 |
| Sod2 | | |
| | HF vs. LF | 0.00002039052 |
| | LF vs. HF+Amp | 0.00000144582 |
| | LF vs. HF+FVT | 0.00017459500 |
| Tff3 | | |
| | LF vs. HF+FVT | 0.02440628000 |
| Timp1 | | |
| | LF vs. HF+Amp+FVT | 0.00899400200 |
| Tjp1 | | |
| | HF vs. HF+Amp | 0.01708157000 |
| Tl1a_Tnfsf15 | | |
| | HF vs. HF+Amp | 0.04826708300 |
| | HF vs. HF+Amp+FVT | 0.02585767930 |
| | HF vs. LF | 0.03749194560 |
| Tlr11 | | |
| | LF vs. HF+Amp+FVT | 0.03096223000 |
| Tlr2 | | |
| | HF vs. HF+Amp | 0.01042556000 |
| | HF vs. HF+Amp+FVT | 0.00305242200 |
| Tlr3 | | |
| | HF vs. HF+Amp | 0.00000490660 |
| | HF vs. HF+Amp+FVT | 0.02508087000 |
| | HF vs. LF | 0.00149313600 |
| | LF vs. HF+FVT | 0.02628533000 |
| Tlr4 | | |
| | LF vs. HF+Amp | 0.01326514000 |
| | LF vs. HF+Amp+FVT | 0.00935164400 |
| Tlr5 | | |
| | HF vs. HF+Amp | 0.00000237901 |
| | HF vs. HF+Amp+FVT | 0.00009386953 |
| | HF vs. LF | 0.00004349276 |
| | LF vs. HF+FVT | 0.00234562700 |
| Tlr9 | | |
| | LF vs. HF+Amp | 0.03403040000 |
| Tnfa | | |
| | LF vs. HF+Amp | 0.00476400900 |
| | LF vs. HF+Amp+FVT | 0.02116594000 |
| Tslp | | |
| | LF vs. HF+Amp+FVT | 0.02108918950 |
| Zbtb16 | | |
| | HF vs. HF+Amp | 0.00169134900 |
| | HF vs. HF+Amp+FVT | 0.00169267100 |

| | | |
|---|---|---|
| | LF vs. HF+Amp+FVT | 0.00005930895 |
| | LF vs. HF+FVT | 0.00000430848 |
| Srebf1a | | |
| | HF vs. LF | 0.00039402150 |
| | LF vs. HF+Amp | 0.00008254950 |
| | LF vs. HF+Amp+FVT | 0.00014998820 |
| | LF vs. HF+FVT | 0.00924378200 |
| Srebp1c | | |
| | HF vs. HF+Amp | 0.01363416000 |
| | HF vs. HF+Amp+FVT | 0.03409613000 |
| | LF vs. HF+Amp | 0.00074078120 |
| | LF vs. HF+Amp+FVT | 0.00219755800 |
| | LF vs. HF+FVT | 0.01898771000 |
| Stat3 | | |
| | HF vs. LF | 0.01857930000 |
| | LF vs. HF+Amp | 0.01485824000 |
| | LF vs. HF+FVT | 0.00556209200 |
| Tlr12 | | |
| | HF vs. LF | 0.00000000643 |
| | LF vs. HF+Amp | 0.00000017364 |
| | LF vs. HF+Amp+FVT | 0.00000015856 |
| | LF vs. HF+FVT | 0.00000014415 |
| Ucp2 | | |
| | HF vs. LF | 0.00092831630 |
| | LF vs. HF+Amp | 0.00066864610 |
| | LF vs. HF+Amp+FVT | 0.00052800160 |
| | LF vs. HF+FVT | 0.00137723080 |
| Usmg5 | | |
| | LF vs. HF+Amp+FVT | 0.01100256000 |
| Tnfa | | |
| | LF vs. HF+Amp | 0.00476400900 |
| | LF vs. HF+FVT | 0.02116594000 |
| Tslp | | |
| | LF vs. HF+FVT | 0.02108918950 |
| Zbtb16 | | |
| | HF vs. HF+Amp | 0.00169134900 |
| | HF vs. HF+FVT | 0.00169267100 |

**TableS4:** Pairwise comparison of α- (Shannon index) and β-diversity (Bray-Curtis dissimilarity) measures of both the bacterial (BC) and viral (VC) community. Statistics were based on Kruskal Wallis for α-diversity and ANOSIM for β-diversity. HF = high-fat, LF = low-fat, FVT = faecal virome transplantation, Amp = Ampicillin.

| | | BC-Bray-Curtis | | BC - Shannon | | VC-Bray-Curtis | | VC - Shannon | |
|---|---|---|---|---|---|---|---|---|---|
| **Group 1** | **Group 2** | **R** | **p-value** | **H** | **p-value** | **R** | **p-value** | **H** | **p-value** |
| HF+Amp | HF+Amp+FVT | 0.780 | 0.001 | 5.835 | 0.016 | 0.875 | 0.001 | 0.176 | 0.674 |
| | HF | 1.000 | 0.001 | 9.926 | 0.002 | 0.904 | 0.001 | 9.926 | 0.002 |
| | LF | 1.000 | 0.001 | 10.500 | 0.001 | 0.922 | 0.001 | 9.276 | 0.002 |
| | HF+FVT | 1.000 | 0.001 | 11.294 | 0.001 | 0.862 | 0.001 | 11.294 | 0.001 |
| HF+Amp+FVT | HF | 0.492 | 0.001 | 0.276 | 0.600 | 0.781 | 0.001 | 11.294 | 0.001 |
| | LF | 0.761 | 0.002 | 10.500 | 0.001 | 0.853 | 0.001 | 11.294 | 0.001 |
| | HF+FVT | 0.598 | 0.002 | 6.353 | 0.012 | 0.910 | 0.001 | 11.294 | 0.001 |
| HF | LF | 0.749 | 0.001 | 7.714 | 0.005 | 0.528 | 0.002 | 0.540 | 0.462 |
| | HF+FVT | 0.362 | 0.003 | 5.835 | 0.016 | 0.502 | 0.002 | 0.276 | 0.600 |
| LF | HF+FVT | 0.741 | 0.001 | 0.054 | 0.817 | 0.773 | 0.001 | 2.824 | 0.093 |

**TableS5:** List of features correlated (Pearson and/or rCCA) to either bacterial abundance or gene expression levels which were further characterised. The suggested metabolite annotation is suggested. Lysophosphatidylcholine = lysoPC.

| Feature ID | Correlated to bacterial abundance | Correlated to gene expressions | Metabolite annotation |
|---|---|---|---|
| 135 | x | - | L-Methionine |
| 707 | x | x | LysoPC(22:4) |
| 713 | x | - | LysoPC(16:0) |
| 774 | x | - | Not identifiable |
| 821 | - | x | Gamma-glutamyltyrosine |
| 850 | x | - | Not identifiable |
| 1189 | x | - | Unknown |
| 1495 | x | - | Unknown |
| 1730 | x | - | PC (16:0/22:6) |
| 2197 | x | - | No biological clusters |
| 2248 | - | x | PC (18:1/O-18:2) |
| 2995 | x | - | No biological clusters |
| 3119 | x | x | LysoPC(22:2) |
| 3510 | x | - | No biological clusters |
| 4014 | x | - | No biological clusters |
| 5650 | x | - | Not identifiable |
| 6103 | x | - | No biological clusters |
| 6261 | x | x | Large PC |
| 7128 | x | - | LysoPC(16:0) |
| 7221 | x | - | LysoPC(18:2) |
| 7440 | x | - | LysoPC(18:0) |

### Supplementary figures

| Images # | VLP count |
|---|---|
| 1 | 308 |
| 2 | 180 |
| 3 | 90 |
| 4 | 80 |
| 5 | 50 |
| 6 | 110 |
| 7 | 60 |
| 8 | 40 |
| 9 | 120 |
| 10 | 30 |
| **Mean VLP count** | 106.8 |
| **Filter area** | 2.84E+08 |
| **Images Area** | 3250 |
| **Images pr. filter** | 8.74E+04 |
| **Virome pr. filter (µL)** | 1 |
| **VLP/mL = (mean VLP) * (Images pr. filter) * 1000 µL** | 4.67E+10 |
| **Diluted FVT virome (VLP/mL)** | 2.00E+10 |

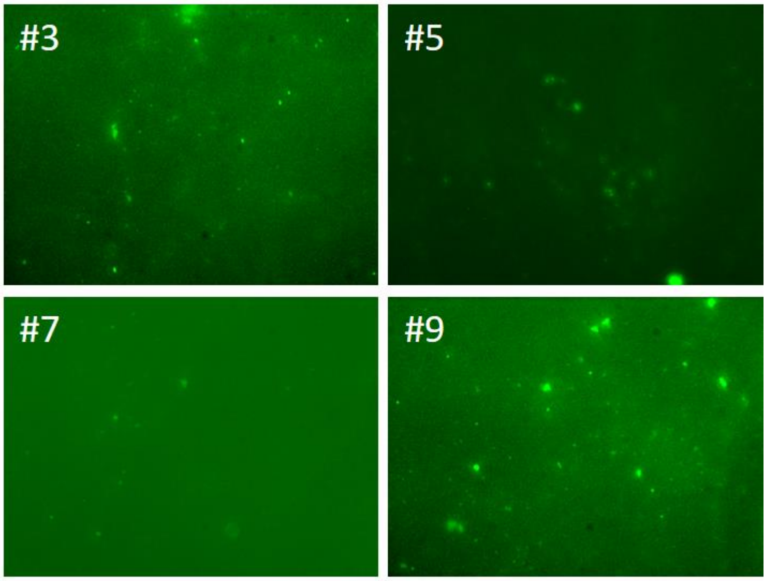


**FigureS1:** Fluorescence microscopy with Zeiss Plan-Neofluar 100x objective of 20x SYBR-gold stained viromes purified through a two-layer CsCl-gradient. Viruses was retained on an Anodiscs® 0.02 µm filter. Numbers inside images refers to image-number in the left panel. VLP = Virus-like particle. FVT = faecal virome transplantation.

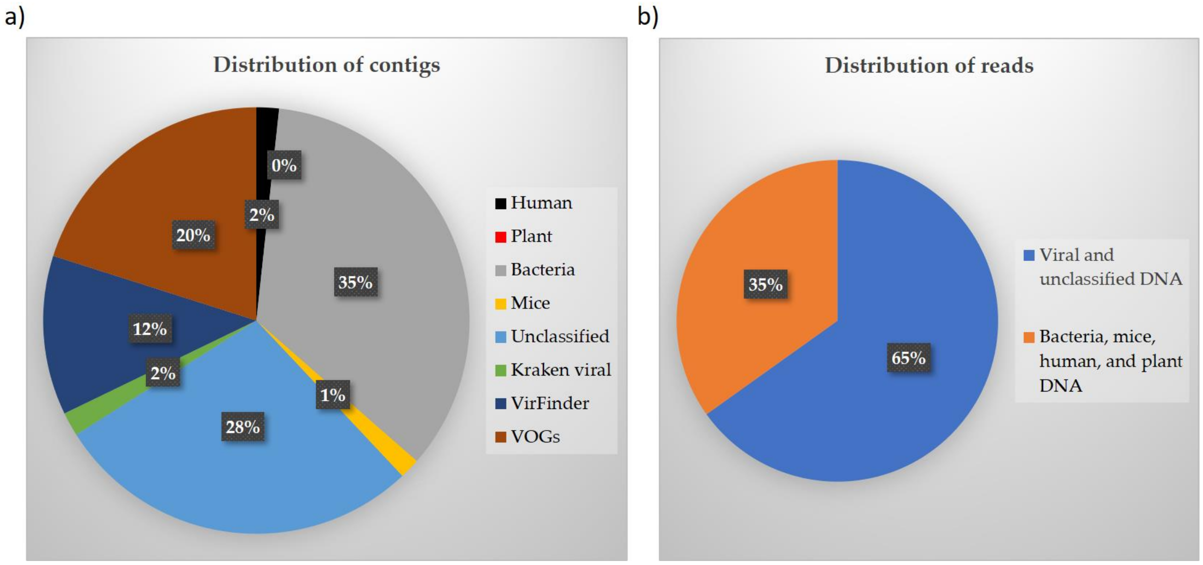


**FigureS2.** Pie plots showing the distribution of the assembled contigs based on origin a) and Illumina NextSeq reads b) of the sequenced metaviromes. Contigs identified by VOGs, Kraken, VirFinder and unclassified DNA constituted the vOTU-table whereas the residuals were removed as contaminations. More than 65% of the Illumina NextSeq reads were used for viral and unclassified contigs (vOTUs).

a)

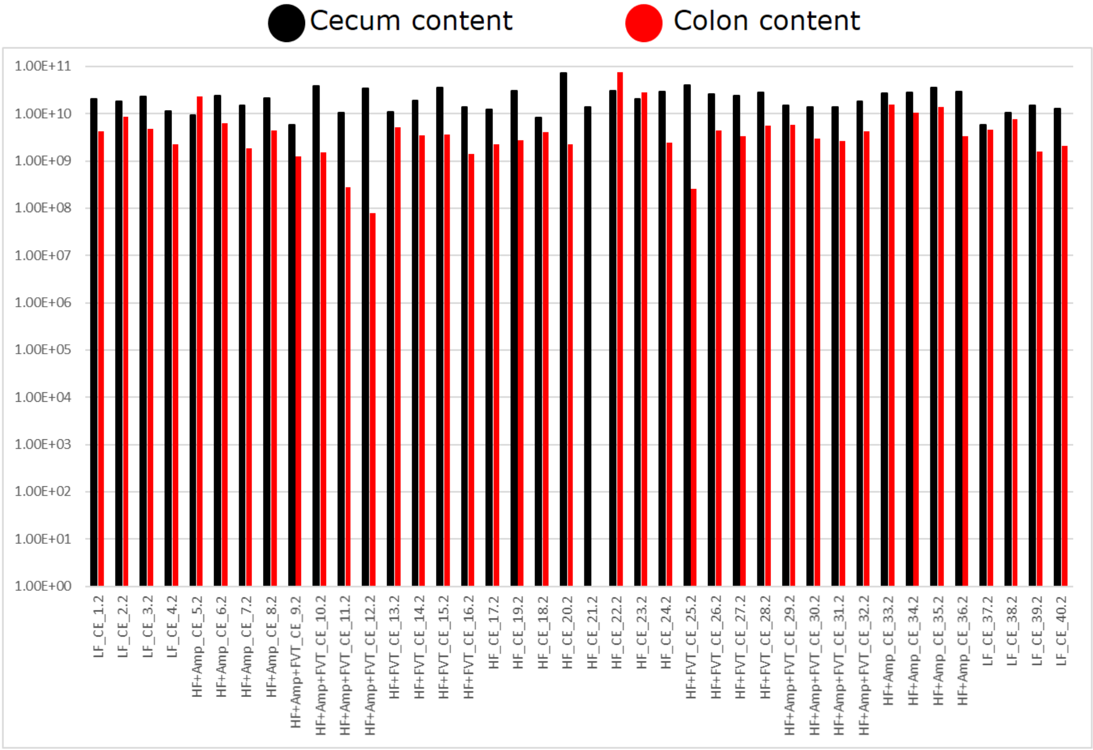


b)

| | Cecum | | | | | Colon LF | | | | |
|---|---|---|---|---|---|---|---|---|---|---|
| **Group** | **LF** | **HF** | **HF+FVT** | **HF+Amp** | **HF+Amp+FVT** | **LF** | **HF** | **HF+FVT** | **HF+Amp** | **HF+Amp+FVT** |
| **Mean 16S rRNA genes/g** | 1.46E+10 | 2.70E+10 | 2.46E+10 | 2.38E+10 | 1.89E+10 | 4.44E+09 | 1.46E+10 | 3.40E+09 | 9.96E+09 | 2.36E+09 |
| **STD** | 5.36E+09 | 1.86E+10 | 9.51E+09 | 8.00E+09 | 1.10E+10 | 2.37E+09 | 2.42E+10 | 1.69E+09 | 6.96E+09 | 1.84E+09 |
| **Sample size (n)** | 8 | 8 | 8 | 8 | 8 | 8 | 7 | 8 | 8 | 8 |
| **Outliers removed** | 0 | 0 | 0 | 0 | 0 | 0 | 0 | 0 | 0 | 0 |

c)

| | | p-value (t-test) | |
|---|---|---|---|
| **Group 1** | **Group 2** | **Cecum** | **Colon** |
| HF | LF | 0.06 | 0.14 |
| HF | HF+FVT | 0.38 | 0.12 |
| HF | HF+Amp | 0.34 | 0.27 |
| HF | HF+Amp+FVT | 0.17 | 0.11 |
| LF | HF+FVT | 0.02 | 0.18 |
| LF | HF+Amp | 0.01 | 0.04 |
| LF | HF+Amp+FVT | 0.19 | 0.04 |
| HF+Amp | HF+Amp+FVT | 0.18 | 0.01 |
| HF+Amp | HF+FVT | 0.43 | 0.02 |
| HF+Amp+FVT | HF+FVT | 0.16 | 0.14 |

**FigureS3:** a) Absolute bacterial abundance determined by qPCR as 16S rRNA gene copies/g (V3 region as described in methods). All samples are included in the bar plot to show sample variations. b) Mean 16S rRNA gene copies/g in cecum and colon samples in the LF, HF, HF+FVT, HF+Amp, HF+Amp+FVT. c) A pairwise t-test of differences between 16S rRNA gene copies/g. HF diet = high-fat, LF = low-fat diet, Amp = ampicillin, FVT = faecal virome transplantation.

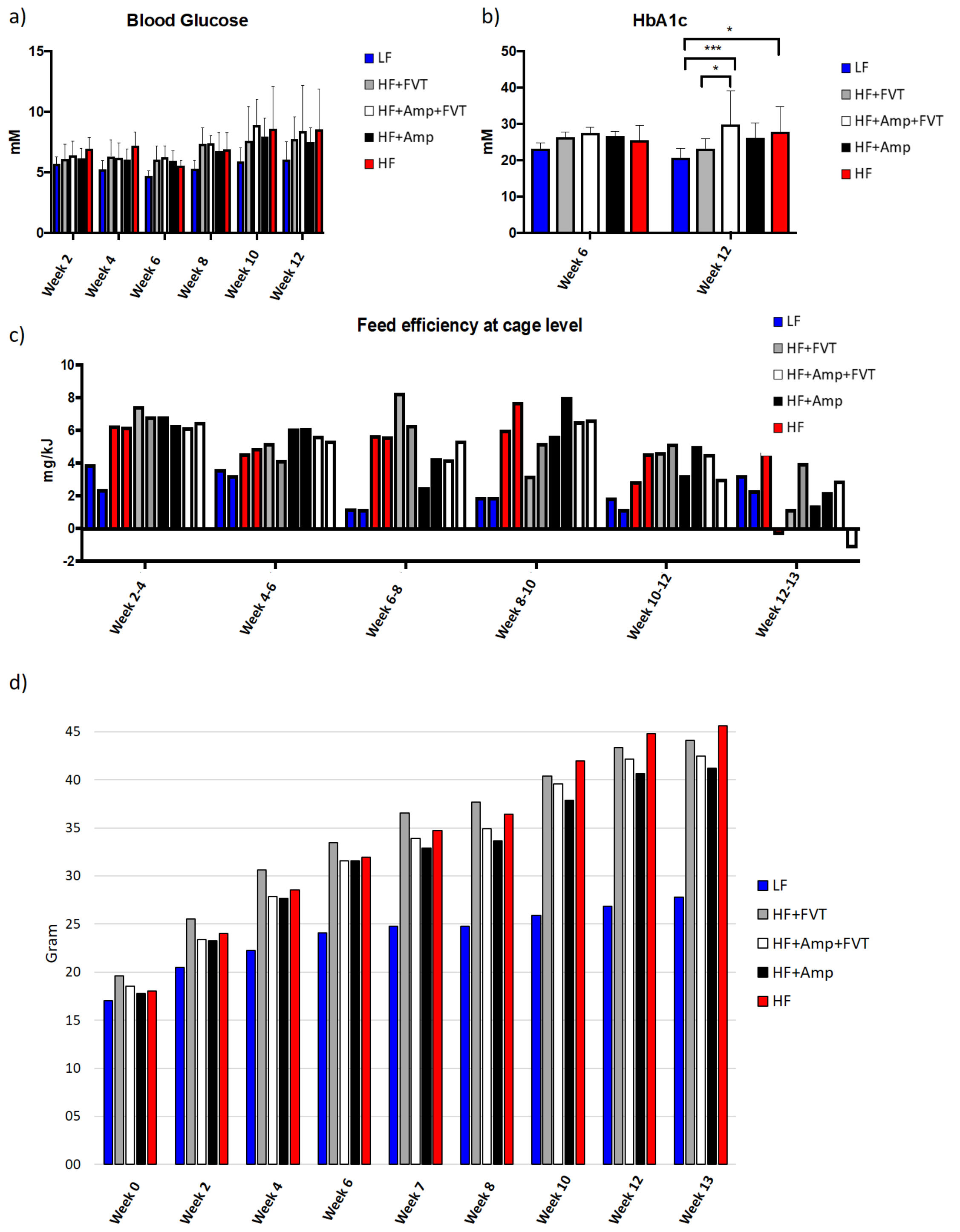


**FigureS4:** a) Non-fasted blood glucose levels measured at several time points (weeks). b) HbA1c (long-term blood sugar) measured pre and post FVT. c) Feed efficiency at cage level measured at several time points evaluated as mg/kJ. d) Development of body mass (gram) in average pr. group measured at several time points. HF diet = high-fat, LF = low-fat diet, Amp = ampicillin, FVT = faecal virome transplantation.

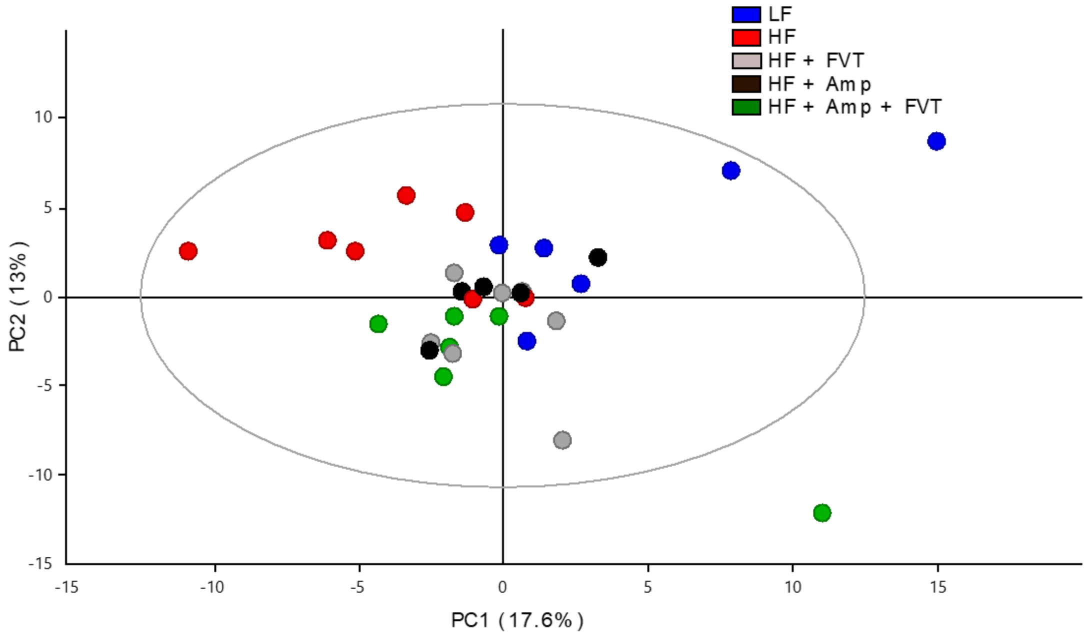


**FigureS5:** PCA scores plot obtained from ESI+ UPLC-MS of plasma from all experimental groups (18 weeks old). (R2=0.47 and Q2=0.10). HF diet = high-fat, LF = low-fat diet, Amp = ampicillin, FVT = faecal virome transplantation.

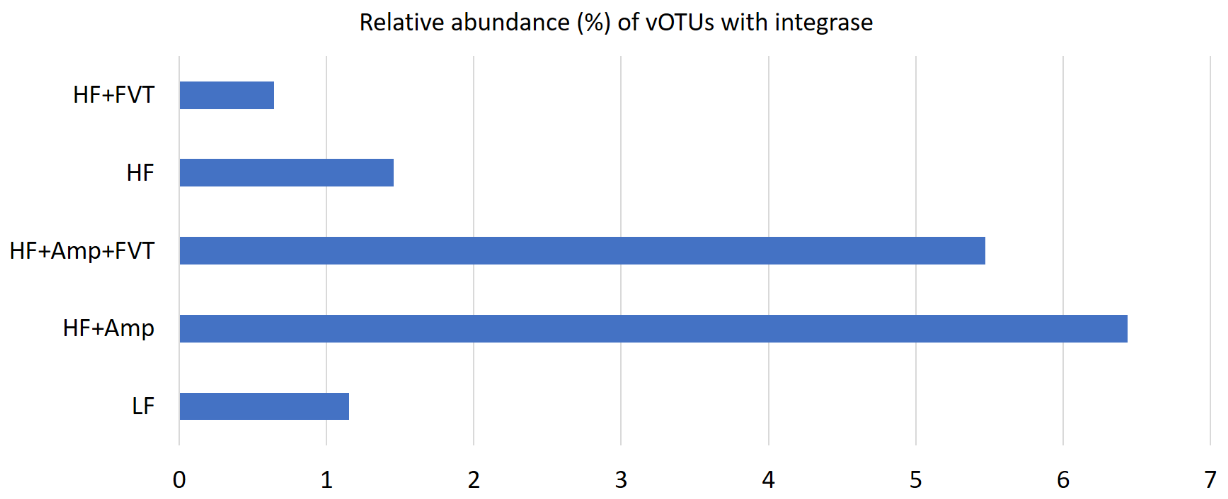


**FigureS6:** Bar plot showing the presence of viral contigs containing integrase genes. BLASTX database was applied to annotate and evaluate the presence of known integrase genes with a minimum E-value = $10^{-3}$ and alignment length at 51 bp amongst vOTU's with a contig size above 3000 bp. HF diet = high-fat, LF = low-fat diet, Amp = ampicillin, FVT = faecal virome transplantation.

See the additional high-resolution PDF FigureS7.

**FigureS7:** Heatmap illustrating regularised Canonical Correlation Analysis (rCCA) with a correlation threshold of r = 0.75, and with component 1:3 included. The components explain 31.6% of the variance of the dataset, and the included bOTUs and vOTUs are in average representing, respectively, 31.5% and 36.2% of the total relative abundance. Only vOTUs ≥ 5000 bp are included. The bOTUs are listed vertically and the vOTUs are listed horizontally.

See the additional high-resolution PDF FigureS8.

**FigureS8:** Heatmap illustrating normalised co-abundance of random forest selected variables with component 1:2 included. The components explain 63.4% of the variance of the dataset, and the included bOTUs and vOTUs are in average representing, respectively, 69.8% and 40.6% of the total relative abundance. Only vOTUs ≥ 5000 bp are included. The bOTUs are listed vertically and the vOTUs are listed horizontally. HF diet = high-fat, LF = low-fat diet, Amp = ampicillin, FVT = faecal virome transplantation.

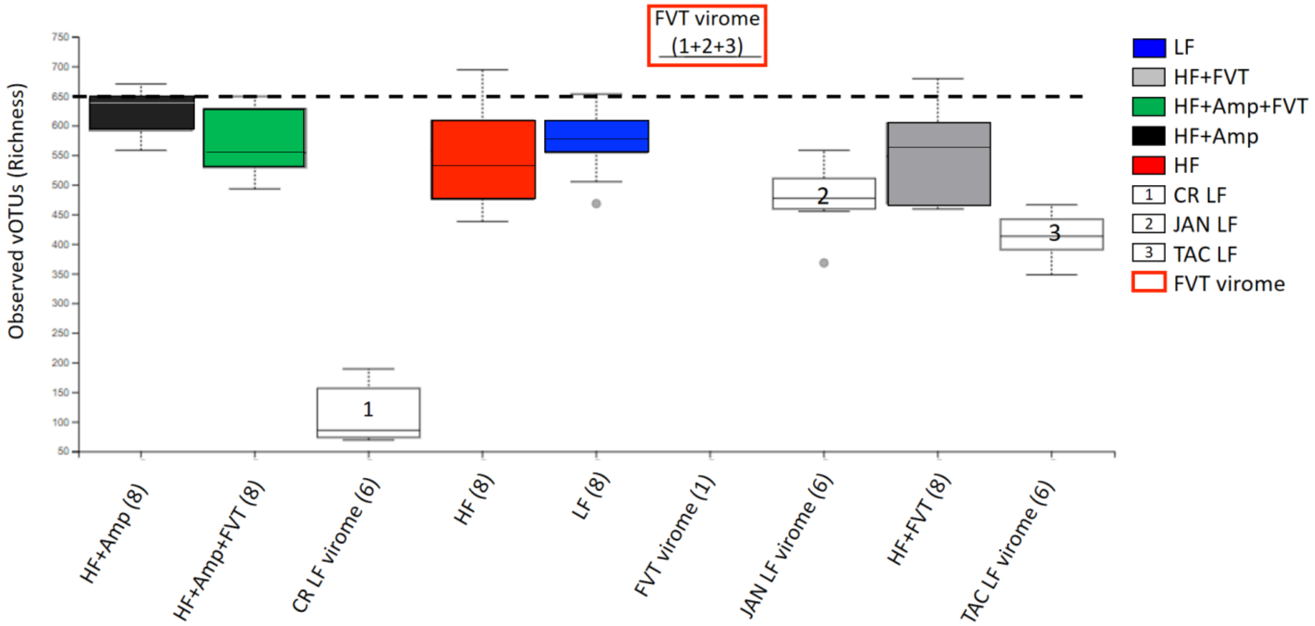


**FigureS9:** Box and whiskers plot showing Caudovirales richness (observed vOTUs) of the experimental groups as well as the FVT virome and its three mixed components (1+2+3). The FVT virome is derived from the cecum of lean mice from three different vendors fed low-fat diet. The mixed FVT virome (1+2+3) is clearly increased compared to the viromes from the individual vendor. Parentheses indicate the number of samples included, and grey dots indicate outliers. HF = high-fat, LF = low-fat diet, Amp = ampicillin, FVT = faecal virome transplantation, 1 = Charles River (CR), 2 = Janvier (JAN), 3 = Taconic (TAC).

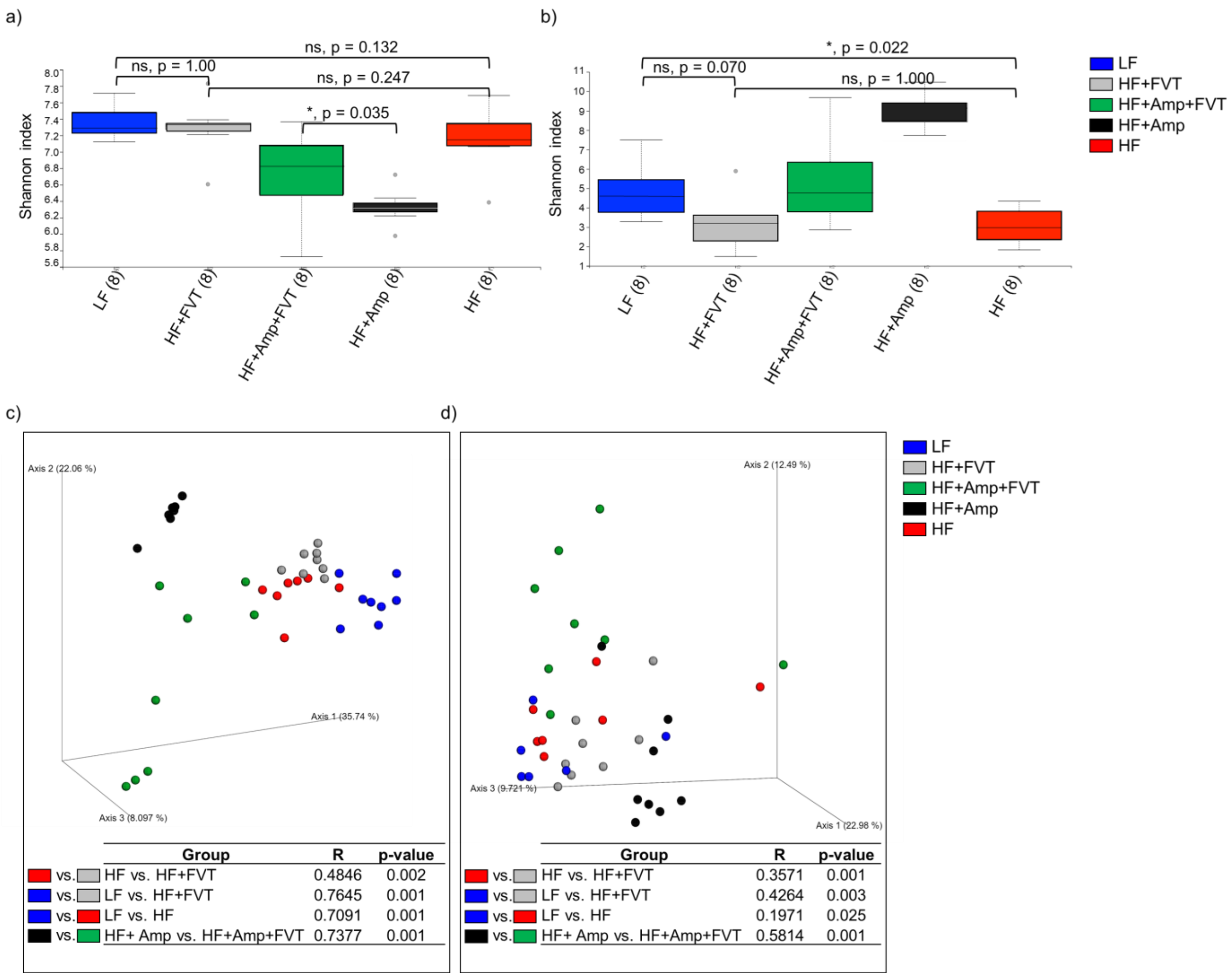


**FigureS10:** Shannon index of the colon a) bacterial and b) viral community at termination (18 weeks old). The parentheses show the number of samples from each group included in the plot and grey dots indicate outliers. Bray Curtis dissimilarity metric PCoA based plots of c) the colon bacterial community and d) viral community at termination (18 weeks old). Tables include ANOSIM of the Bray Curtis dissimilarity to show the effect of the FMT on the GM composition. HF = high-fat, LF = low-fat diet, Amp = ampicillin, FVT = faecal virome transplantation
